# Transit mortality and the efficiency of *Tamarixia radiata* releases against *Diaphorina citri* in metapopulation networks

**DOI:** 10.64898/2026.11.12.766282

**Authors:** Sebastián Rodríguez-Falcón, Heliana Arias-Castro, Javier Galeano, Luciano Stucchi

**Affiliations:** Pontificia Universidad Católica del Perú, Lima, Peru; Universidad del Valle, Calle 13 100-00, Cali 760032, Colombia; Grupo Interdisciplinar de Sistemas Complejos (GISC), Universidad Politécnica de Madrid, Avda. Puerta de Hierro 2-4, 28040 Madrid, Spain; Universidad del Pacífico, Lima, Peru

**Author notes:** Author for correspondence: Luciano Stucchi.

**Keywords:** *Diaphorina citri*, *Tamarixia radiata*, augmentative biological control, metapopulation networks, transit mortality, master stability function

## Abstract

Augmentative releases of the parasitoid *Tamarixia radiata* are a main tool against *Diaphorina citri*, the vector of huanglongbing in citrus, but release models usually treat orchards as a single patch. We extended a four-compartment model of psyllid, flush, parasitoid and plant vigour with impulsive releases to networks of eight patches coupled by diffusion of psyllids and parasitoids, with optional transit mortality, and compared uniform releases with releases targeted at the outbreak patch. Without transit mortality the amount released dominated the outcome. Uniform releases reduced the network pest load by about 30% and targeted releases by 2.5–4.5%, efficiencies per parasitoid were similar, and topology changed the pest load by less than 2%, mainly through the degree of the outbreak patch. The master stability function ranked the networks by spatial coherence but did not account for its level over an outbreak. Parasitoid transit mortality created an intermediate dispersal rate that maximised the efficiency of targeted releases without minimising the pest load. This optimum lay where transit losses pushed the resident parasitoid towards collapse, so that the pest load without releases rose sharply. A linear invasion argument explains why it scales with the mean degree and the mean edge mortality.

## 1. Introduction

Huanglongbing (HLB), or citrus greening, is widely regarded as the most destructive disease affecting citrus worldwide, caused by the phloem-restricted bacterium *Candidatus* Liberibacter spp. and transmitted primarily by the Asian citrus psyllid *Diaphorina citri* [1, 2]. Because no curative treatment exists for the pathogen once a tree is infected, management relies almost entirely on suppressing the vector, and the ectoparasitoid *Tamarixia radiata* is the natural enemy most widely deployed in classical and augmentative biological control programmes against *D. citri* [3, 4]. We recently introduced, and analysed at the scale of a single orchard, a four-compartment model that couples the psyllid, the flush and vigour dynamics of its host plant, and *T. radiata* under periodic augmentative releases [5]. That single-patch model showed that the flush–vigour feedback generates a sustained, endogenous oscillation, that periodic releases interact with this oscillation rather than entrain it, and that sustained releases outperform a single introduction. It also derived an invasion-exponent criterion that separates release schedules that merely suppress the pest from those predicted to drive it towards local eradication.

That analysis, however, treated the agroecosystem as a single, well-mixed patch, an idealisation shared by most of the theoretical literature on augmentative releases, from the early spatial simulation models developed for boll-weevil parasitoids [6] to recent optimal-release studies for egg-parasitoid programmes against stem borers [7]. Citrus is grown, and HLB spreads, across orchards, groves and regions connected by psyllid flight, wind-assisted dispersal and the movement of nursery material, so that a release programme designed for one patch is in practice deployed on a network of patches that continuously exchange both the pest and its natural enemy. In host–parasitoid systems, the spatial arrangement of refuges and the connectivity of the landscape shape persistence and the suppression that can be achieved [8]. How precisely a release is placed relative to the host patch measurably changes realised parasitism [9]. Despite this, explicit network treatments of augmentative release strategies remain scarce. As recently noted for the sterile insect technique, metapopulation models are commonplace in epidemiology but their application to pest control, and in particular to how a release strategy should be chosen for a given dispersal network, is still limited [10].

Representing a population as patches coupled by dispersal along the edges of a graph is the standard metapopulation formulation in epidemiology [11, 12]. In that setting, the degree of the node where a perturbation starts governs whether it stays confined or invades the whole network, and highly connected nodes can act as reservoirs that sustain and re-seed the rest of it [12]. Network structure also matters for purely ecological dynamics. Embedding a Lotka–Volterra community in a network of diffusively coupled patches, rather than treating it as an isolated or merely migration-open system, changes which macroecological patterns the model can reproduce [13]. Moving from a single patch to a network can therefore change which outcomes are reachable. For an augmentative pest–parasitoid system we ask whether the degree of the outbreak patch controls the outcome, as the reservoir-node intuition from epidemic spreading suggests, whether the resulting spatial coherence follows the linear stability of the synchronous state given by the master stability function [14] or the nonlinear, finite-time dynamics of an outbreak, and how dispersal and mortality in transit between patches interact with the choice between releasing uniformly across the network and targeting the outbreak patch alone. These questions have clear analogues in network epidemiology but have not, to our knowledge, been examined together for release-driven biological control.

Here we extend the single-patch model of Rodríguez-Falcón et al. [5] to an explicit network of *n* patches, coupling psyllids and parasitoids by diffusion along the edges of seven graph families that span regular, random, and scale-free topologies, while flush and plant vigour remain local to each patch, as in the original formulation. One patch is seeded with an outbreak, and augmentative releases of *T. radiata* are applied either uniformly across the network or targeted at the outbreak patch alone. This choice is the network analogue of the population-wide and targeted interventions long contrasted in epidemic control on networks [11, 12]. We quantify the outcome using the network-wide pest load and the pest load avoided per parasitoid released. This efficiency measure lets us compare strategies using different numbers of parasitoids, and we complement the simulations with a master-stability analysis of the uniform-release synchronous state.

## 2. Methods

### (a) Local model

Each patch (node) of the network carries four state variables, namely the density of the Asian citrus psyllid *Diaphorina citri* (*x*_1_), the availability of young shoots or flush on which the psyllid feeds and breeds (*x*_2_), the density of the ectoparasitoid *Tamarixia radiata* (*x*_3_), and an index of plant vigour (*x*_4_). The psyllid is the vector of the bacteria associated with huanglongbing [15, 16], and *T. radiata* is the parasitoid most widely used for its augmentative control [4]. All quantities, including time, are in arbitrary units (a.u.). The within-patch dynamics are those of the single-node model [5], which we write as

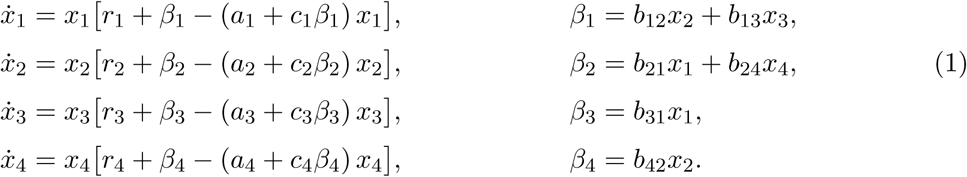

Here *r_i_* is the per capita growth rate of population *x_i_* in the absence of interactions, *b_ij_* is the effect of population *x_j_* on the per capita growth of *x_i_*, and the crowding term *a_i_* + *c_i_β_i_* lets self-limitation depend on the interaction input *β_i_*. The interaction web is shown in figure 1a. Flush benefits the psyllid (*b*_12_ *>* 0) and is damaged by it (*b*_21_ *<* 0). Parasitism lowers psyllid growth (*b*_13_ *<* 0) and sustains the parasitoid (*b*_31_ *>* 0). Vigour promotes flushing (*b*_24_ *>* 0) and is drawn down by it (*b*_42_ *<* 0). Only vigour has a positive intrinsic rate. Parameter values are listed in table 1 and were used unchanged in every experiment except for the sensitivity analysis.

**Figure 1:**
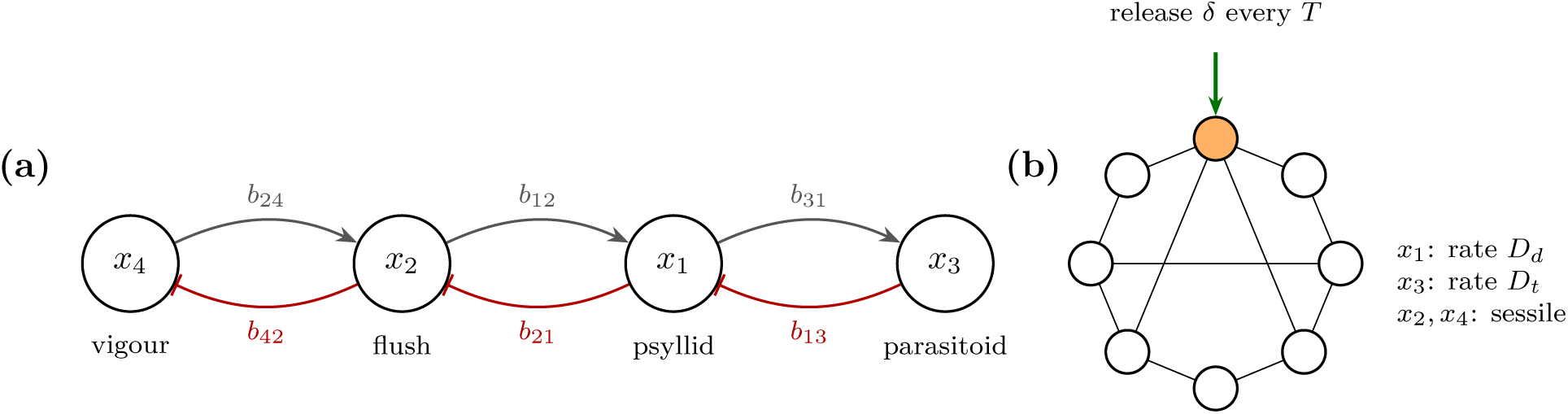
Model structure. (*a*) Interaction web inside one patch. Arrows are positive effects on per capita growth and bars are negative effects. (*b*) Patches are the nodes of a graph. Psyllids and parasitoids disperse along edges, flush and vigour do not. The shaded node is the outbreak patch, which receives all releases under the targeted strategy. The graph drawn is illustrative only.

**Table 1:** Parameters of the local model (equation (1)). The same crowding coefficients *a_i_* = 5.55 × 10*^−^*^4^ and *c_i_* = 5 × 10*^−^*^3^ are used for all four variables.

| variable | intrinsic rate | interaction coefficients | crowding |
|---|---|---|---|
| *x*<sub>1</sub> psyllid | *r*<sub>1</sub> = −0.20 | *b*<sub>12</sub> = 0.040, *b*<sub>13</sub> = −0.010 | *a*<sub>1</sub>, *c*<sub>1</sub> |
| *x*<sub>2</sub> flush | *r*<sub>2</sub> = −0.72 | *b*<sub>21</sub> = −0.010, *b*<sub>24</sub> = 0.054 | *a*<sub>2</sub>, *c*<sub>2</sub> |
| *x*<sub>3</sub> parasitoid | *r*<sub>3</sub> = −0.30 | *b*<sub>31</sub> = 0.010 | *a*<sub>3</sub>, *c*<sub>3</sub> |
| *x*<sub>4</sub> plant vigour | *r*<sub>4</sub> = 0.72 | *b*<sub>42</sub> = −0.072 | *a*<sub>4</sub>, *c*<sub>4</sub> |

The crowding terms *a_i_* + *c_i_β_i_* are not guaranteed to be positive. With the values of table 1, *a*_4_ + *c*_4_*b*_42_*x*_2_ *<* 0 whenever *x*_2_ *>* 1.54, which held at every sampled time of the single-node trajectories we checked (initial state (50, 10, 15, 30), with and without releases). Boundedness of *x*_4_ therefore rests on the term *b*_42_*x*_2_ in its growth rate and not on self-limitation. The psyllid crowding term *a*_1_ + *c*_1_*β*_1_ stayed positive under the reference release schedule but became negative during 15% of the horizon under the most intense schedule tested (*δ* = 50, *T* = 5).

### (b) Network model and parasitoid releases

Patches are the *n* = 8 nodes of an undirected, unweighted graph with adjacency matrix *A* and Laplacian *L* = *K* − *A*, where *K* is the diagonal matrix of node degrees *k_i_*. Psyllids and parasitoids move between neighbouring patches by linear diffusion with rates *D_d_* and *D_t_*, while flush and vigour belong to the host plants and do not move. Individuals may die while moving between patches. We write *m_d_* and *m_t_* for the probabilities that a psyllid or a parasitoid leaving a patch dies before reaching the neighbouring one. Writing *f_k_* for the right-hand side of the *k*th line of equation (1), the network model is

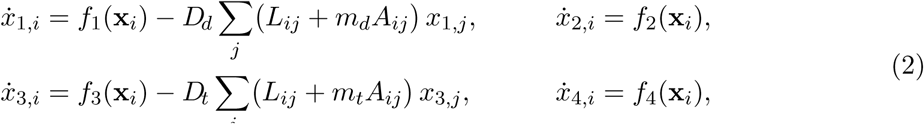

for *i* = 1*, . . ., n*. Emigration from patch *i* occurs at rate *Dk_i_x_i_* whatever the value of *m*, and immigration from each neighbour *j* is reduced from *Dx_j_* to (1 − *m*)*Dx_j_*. Summing over patches gives 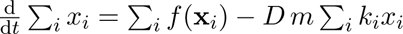 for each dispersing variable. Transit mortality therefore removes individuals at a rate proportional to the degree of the patch they leave. With *m_d_* = *m_t_* = 0, which is the case in Experiments 2, 3 and 6–9, the rows of *L* sum to zero and dispersal conserves the network totals of *x*_1_ and *x*_3_. In Experiment 11 (section 2(e)) the scalar *m_t_* is replaced by an edge-specific probability *m_ij_*, so that the term *m_t_A_ij_* becomes *m_ij_A_ij_*.

Augmentative control is represented as instantaneous releases of *δ* parasitoids at the times *t_k_* = *kT*, *k* = 1, 2*, . . .*, into every node of a release set R,

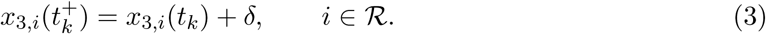

The first release occurs at *t* = *T*, not at *t* = 0. Releases are applied only for *t_k_ < t*_max_, so a simulation of length *t*_max_ receives *N* (*T*) = ⌈*t*_max_*/T* ⌉ − 1 release events. We compared three strategies. *No release* sets *δ* = 0. *Uniform* releases into all *n* patches. *Targeted at source* releases only into the outbreak patch defined below. For a given schedule, a uniform programme therefore uses *n* times as many parasitoids as a targeted one, and the total released is *Q* = *δ N* (*T*) |R|. Impulsive release models of this kind are standard in the theory of integrated pest management [17].

We used seven graph families on *n* = 8 nodes. Four are deterministic: the path, the cycle, the star with the outbreak patch at the hub, and the complete graph. Three are stochastic: Gilbert random graphs *G*(*n, p*) with *p* = 0.4 [18], Watts–Strogatz small-world graphs obtained from a ring in which each node is joined to its four nearest neighbours by rewiring each edge with probability 0.2 [19], and Barabási–Albert graphs with two edges per new node [20]. Graphs were generated with NetworkX [21] using seed 1 unless stated otherwise. Experiment 3 drew ten independent instances (seeds 1–10) of each stochastic family, giving 34 distinct graphs in total. All 34 were connected, and their algebraic connectivity *λ*_2_(*L*) [22] ranged from 0.152 (path) to 8 (complete graph). The outbreak patch is node 1 in every graph. Its degree, which we call the source degree, varied between 1 and 7 across the 34 graphs.

### (c) Initial conditions and numerical solution

All network experiments simulate an outbreak over *t*_max_ = 100 a.u., unless stated otherwise. We used two initial conditions. In IC-A, which is the condition of Experiments 2, 3, 6 and 7, the outbreak patch starts with *x*_1_ = 200 and the other seven patches with *x*_1_ = 20, so the outbreak arises on a background infestation. In IC-B, a point introduction, the outbreak patch starts with *x*_1_ = 200 and the other patches with *x*_1_ = 0. Because *x*_1_ = 0 is invariant for the local dynamics, under IC-B the psyllid reaches the other patches only by dispersal, and *D_d_* = 0 is excluded. In both conditions every patch starts with *x*_2_ = 10, *x*_3_ = 15 and *x*_4_ = 30.

Equations (2) were integrated with the explicit Runge–Kutta method of order 8(5,3) of Dormand and Prince as implemented in SciPy [23, 24], with relative and absolute tolerances 10*^−^*^10^ and 10*^−^*^12^. The parameter sweeps of Experiments 8–11 used tolerances of 10^−8^ and 10^−10^. On four test configurations spanning *D_t_* from 10*^−^*^3^ to 1, this changed the efficiencies defined below by less than 10*^−^*^8^ in relative terms and reduced computing time by about 40%. Releases were handled exactly by integrating the smooth system between consecutive release times and applying the jump (3) to the end state of each piece. The resulting solution is right-continuous, so its value at a release time is the post-release state. Time integrals were computed from the dense output of the integrator by eight-point Gauss–Legendre quadrature on every internal step, which is exact for the degree-7 interpolant. As a consistency check, we set *D_d_* = *D_t_* = 0 and *δ* = 0 on a three-node path with identical initial states. The maximum absolute deviation of each node from an independently coded single-patch solution over *t* ∈ [0, 100] was 1.8 × 10*^−^*^13^ for *x*_1_ and 6.0 × 10*^−^*^14^ for *x*_3_, and the three nodes were identical to machine precision.

The version of the code with transit mortality passed four further checks before it was used. (i) With *m_d_* = *m_t_* = 0 it reproduced the network AUC of the 336 runs of Experiment 2 with a maximum relative error of 4.1 × 10*^−^*^16^. (ii) With the local dynamics switched off, the total of each dispersing variable decayed at the rate *D m* Σ*_i_ k_i_x_i_* derived above, with relative errors below 1.3 × 10*^−^*^15^ in the right-hand side and 4.5 × 10*^−^*^11^ in the integrated totals, over five graphs and three pairs (*D, m*). (iii) With *D* = 0, every value of *m* gave results identical to *m* = 0. (iv) With *m* = 0 the star and the complete graph gave the same AUC to within 10*^−^*^14^ under both initial conditions and all strategies, as expected (section 3(c)). An edge-specific matrix *m_ij_* with all entries equal to *m* also reproduced the scalar case.

The model code is a Python port of an original Mathematica implementation and requires NumPy ≥ 1.24, SciPy ≥ 1.10 and NetworkX ≥ 3.0. The post hoc analyses were run with NumPy 2.4.4, SciPy 1.17.1 and NetworkX 3.6.1. Experiments 8–11 and the checks of the transit-mortality code were run with NumPy 2.5.3, SciPy 1.18.1 and NetworkX 3.6.1. All software is available in our repository [25].

### (d) Response variables

The primary response is the network pest load, the area under the psyllid curves summed over patches,

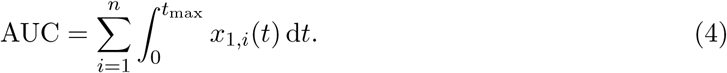

We also report the mean psyllid density per patch over the second half of the horizon, [*t*_max_*/*2*, t*_max_]. Spatial heterogeneity was measured by the coefficient of variation across patches of the patch mean of *x*_1_ over [*t*_0_*, t*_max_], with *t*_0_ = *t*_max_ − (*t*_max_ mod *T*) − *T*, which is [75, 100] for *T* = 15. Spatial coherence was measured by the synchrony index, the mean Pearson correlation between the psyllid time series of all pairs of patches sampled at 401 equally spaced times on [0*, t*_max_]. The synchrony index includes the initial transient.

Strategies that use different numbers of parasitoids were compared through their efficiency. The AUC avoided is ΔAUC = AUC_0_ − AUC, where AUC_0_ is the no-release run on the same graph and with the same dispersal rates. The efficiency of a schedule is ΔAUC*/Q*, the pest load avoided per parasitoid released.

In Experiments 8–11 the three strategies were always solved together, with identical parameters, and AUC_0_ is the no-release run with the same graph, initial condition, dispersal rates, transit mortalities and *r*_3_. We write *E_T_* and *E_U_*for the efficiencies of targeted and uniform releases, with *Q_T_* = *δ N* (*T*) and *Q_U_* = *n δ N* (*T*), and use *E_T_ /E_U_* to indicate when concentrating the releases at the outbreak patch pays off. Because AUC_0_ itself depends on every parameter that is varied, an efficiency can rise either because releases suppress the pest more or because the pest load without releases is larger.

To ask whether *E_T_* depends on a single combination of parameters, we defined the lifetime of a parasitoid without hosts, *τ_t_* = 1*/*|*r*_3_|, and two candidate time scales for the psyllid, *τ_d_* = 1*/*|*r*_1_| = 5 a.u. and the characteristic period of the isolated patch, *T*_0_ ≈ 7.8 a.u. (section 3(a)). The relative dispersal

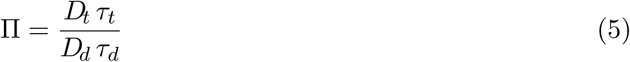

compares how far each species disperses during its own time scale and does not depend on degree. The reach of the released parasitoids,

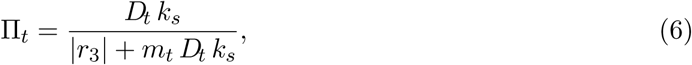

is the expected number of departures from the outbreak patch, of degree *k_s_*, during the lifetime of a parasitoid there. With transit mortality the total loss rate at that patch is |*r*_3_| + *m_t_D_t_k_s_*, so Π*_t_* is bounded by 1*/m_t_* however large *D_t_*becomes.

Associations between network descriptors and outcomes across the 34 graphs of Experiment 3 were measured with Spearman rank correlations at *D_d_* = *D_t_* = 0.03. The descriptors were the source degree, *λ*_2_ and the mean degree. The source degree was recovered by regenerating each graph with the same generator and seed. The recomputed *λ*_2_ matched the stored values to within 4 × 10*^−^*^15^, which confirms that the regenerated graphs are the ones simulated.

### (e) Simulation experiments

The experiments are summarised in table 2. Experiment 1, which validated the Python port of the original code, is not reported here; it is included in our repository [25]. Experiment 2 crossed the seven topologies (seed 1), four values of each dispersal rate, *D_d_, D_t_* ∈ {0, 0.01, 0.03, 0.1}, and the three strategies, with the reference schedule *δ* = 35, *T* = 15. Experiment 3 repeated this design on all 34 graph instances to quantify the variation among realisations of the stochastic families. Experiment 6 fixed *D_d_* = *D_t_* = 0.03 and crossed four topologies (path, star, complete, and random) with a 5 × 5 grid of release sizes and periods and the two release strategies. Experiment 7 is the sensitivity analysis, and Experiments 8–11 study relative dispersal and transit mortality. Both are described below.

**Table 2:** Design of the simulation experiments. Experiments 4 and 5 are single-patch or linearised analyses and are described in section 2(f). Unless stated otherwise, *δ* = 35, *T* = 15, *t*_max_ = 100, *r*_3_ = −0.3 and *m_d_* = *m_t_*= 0. In Experiments 8–11, each combination of parameters was solved under the three strategies (two in Experiment 11, where uniform releases were not needed), and the run counts include them.

| experiment | factors and levels | runs |
|---|---|---|
| 2 | 7 topologies × 4 *D<sub>d</sub>* × 4 *D<sub>t</sub>* × 3 strategies; *δ* = 35, *T* = 15 | 336 |
| 3 | 34 graphs (stochastic families, 10 seeds) × 4 *D<sub>d</sub>* × 4 *D<sub>t</sub>* × 3 strategies | 1632 |
| 6 | 4 topologies × *δ* ∈ {10, . . . , 50} × *T* ∈ {5, 10, 15, 20, 30} × 2 strategies; *D<sub>d</sub>* = *D<sub>t</sub>* = 0.03 | 200 + 4 |
| 7 | 18 parameters × {0.9, 1.1}; complete graph, uniform, *δ* = 35, *T* = 15, *D<sub>d</sub>* = *D<sub>t</sub>* = 0.03 | 36 + 1 |
| 8 | 9 *D<sub>d</sub>* × 9 *D<sub>t</sub>* (log-spaced, 10<sup>−3</sup>–1) × 3 *r*<sub>3</sub> × 7 focal graphs × 2 ICs × 3 strategies; *m<sub>d</sub>* = *m<sub>t</sub>* = 0 | 10 206 |
| 9 | 25 *D<sub>t</sub>* × 3 *D<sub>d</sub>* × 7 focal graphs × 2 ICs × 2 *t*<sub>max</sub> × 3 strategies; *m<sub>d</sub>* = *m<sub>t</sub>* = 0. Check at *D<sub>d</sub>* = 0: 26 *D<sub>t</sub>* × 7 graphs × 2 *t*<sub>max</sub> × 3 strategies, IC-A | 6300 + 1092 |
| 10a | 5 *m<sub>t</sub>* × 13 *D<sub>t</sub>* × 3 *D<sub>d</sub>* × 8 graphs (7 focal + complete) × 2 ICs × 2 *r*<sub>3</sub> × 3 strategies; *m<sub>d</sub>* = 0 | 18 720 |
| 10b | 2 *m<sub>d</sub>* × 4 *m<sub>t</sub>* × 7 *D<sub>t</sub>* (≥ 0.03) × 3 *D<sub>d</sub>* × 8 graphs × 3 strategies; IC-B, *r*<sub>3</sub> = −0.3 | 4032 |
| 11 | 10 random geometric graphs × 3 *ℓ* × 15 *D<sub>t</sub>* (0.03–3) × 2 strategies; *m<sub>ij</sub>* = 1 − *e*<sup>−*d<sub>ij</sub>*/*ℓ*</sup>, IC-B, *D<sub>d</sub>* = 0.03 | 900 |

### Experiment 7, sensitivity analysis

Local one-at-a-time sensitivity was evaluated on the complete graph with *D_d_* = *D_t_*= 0.03, uniform releases, *δ* = 35 and *T* = 15. Each of the 18 parameters of table 1 was multiplied by 0.9 and by 1.1 while the others were held at their reference values, and we recorded the percentage change in network AUC relative to the reference run. Parameters were ranked by the larger of the two absolute changes. Because the perturbation is multiplicative, the factor 1.1 increases the magnitude of negative parameters. For example, it makes *r*_3_ more negative, which corresponds to higher parasitoid mortality. A one-at-a-time design explores only a small neighbourhood of the reference point and ignores interactions between parameters [26], so these results describe local behaviour only.

### Focal graphs

To vary the degree of the outbreak patch independently of the graph family, Experiments 8–10 used seven graphs on *n* = 8 nodes whose outbreak patch has degree *k_s_* = 1*, . . .,* 7. They are the path, with the outbreak at one end (*k_s_* = 1), the cycle (*k_s_* = 2), three Gilbert graphs of Experiment 3 (seeds 9, 3 and 4, with *k_s_* = 3, 4 and 5), a Barabási–Albert graph of Experiment 3 (seed 2, *k_s_* = 6) and the star (*k_s_* = 7). For each degree we took the first seed between 1 and 10 that produced it. No Gilbert instance had an outbreak patch of degree 6, which is why a Barabási–Albert graph fills that slot. The mean degrees ⟨*k*⟩ of the seven graphs are 1.75, 2, 3, 2.5, 3.25, 3 and 1.75. Experiment 10 added the complete graph (*k_s_* = ⟨*k*⟩ = 7). With *m* = 0 it is equivalent to the star (section 3(c)), but with *m_t_ >* 0 it is not.

### Experiment 8, relative dispersal without transit cost

*D_d_* and *D_t_* each took nine log-spaced values between 10*^−^*^3^ and 1, and *r*_3_ ∈ {−0.2, −0.3, −0.45}, so that *τ_t_*varies and Π is not just *D_t_/D_d_*. We crossed these with the seven focal graphs, both initial conditions and the three strategies. A pilot of 243 runs on the complete graph (IC-B, *r*_3_ = −0.3) preceded the full design. We asked whether *E_T_*collapses onto a single candidate variable. The candidates were *D_t_*, *D_t_/D_d_*, Π with each choice of *τ_d_*, and the pair (Π, Π*_t_*). For each candidate we fitted a cubic polynomial in log_10_ of the candidate (a bivariate cubic for the pair) and computed *R*^2^ on held-out data, leaving out one value of *r*_3_ at a time (three folds) or one graph at a time (seven folds). Held-out values outside the training range were clamped to it, so the fitted curve was extended as a constant rather than extrapolated. We used the smaller of the two cross-validated *R*^2^. A candidate was said to collapse the data if this value was at least 0.9 and exceeded that of *D_t_/D_d_* by at least 0.1. As an upper reference we also fitted a free bivariate cubic in (log *D_d_,* log *D_t_*), which bounds what any function of the two dispersal rates can explain. Because neither *r*_1_ nor *T*_0_ was varied, the two versions of Π differ by the constant factor *T*_0_|*r*_1_|, and their fits are identical.

### Experiment 9, parasitoid dispersal rate without transit cost

*D_t_* took 25 log-spaced values between 10*^−^*^3^ and 1, with *D_d_* ∈ {0.01, 0.03, 0.1}, the seven focal graphs, both initial conditions and horizons *t*_max_ = 100 and 300 a.u. For each curve *E_T_* (*D_t_*) we located the grid maximum and, when it was interior, refined its position *D_i_*\* with a parabola in log_10_ *D_t_* through the maximum and its two neighbours. The prominence of the maximum is the relative drop 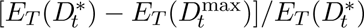 to the upper end of the grid, *D_t_*^max^, which is 1 in Experiments 9 and 10 and 3 in Experiment 11. An interior maximum was considered real if its prominence exceeded 5% under both horizons. A separate set of runs with *D_d_* = 0 and IC-A, including *D_t_* = 0, examined on a fine grid the decrease of *E_T_* with *D_t_* that Experiment 2 showed at *D_d_* = 0.

### Experiment 10, transit mortality

In Experiment 10a only parasitoids paid a cost, with *m_t_* ∈ {0, 0.05, 0.1, 0.2, 0.4} and *m_d_*= 0. *D_t_* took 13 log-spaced values between 10*^−^*^3^ and 1, with *D_d_*∈ {0.01, 0.03, 0.1}, the eight graphs, IC-B as the main case and IC-A as control, and *r*_3_ ∈ {−0.3, −0.45}. A pilot of 585 runs on the complete graph (IC-B, *r*_3_ = −0.3) preceded it. Optima were located as in Experiment 9. Because only the horizon *t*_max_ = 100 was simulated, a maximum was considered real if it was interior with prominence above 5% at that horizon alone, a weaker requirement than in Experiment 9. To test whether the optimum lies where transit losses equal natural mortality, we computed the ratio 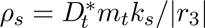 and, replacing the source degree by the mean degree, 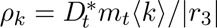. We also recorded the pest AUC with and without releases and the parasitoid AUC without releases at D*_t_*\*, to identify what drives the optimum, and the Pearson correlation across the seven focal graphs between *k_s_* and the AUC under targeted releases. For each curve of the pest AUC under targeted releases against *D_t_* we recorded whether it had an interior minimum and its depth relative to the lower of the two ends of the curve. Under IC-B, for every combination of graph, *D_d_*and *r*_3_ with real maxima at two or more values of *m_t_*, we fitted the slope of log_10_ D*_t_*\* against log_10_ *m_t_*by least squares. Experiment 10b added psyllid transit mortality *m_d_* ∈ {0.1, 0.2} on a reduced grid around the optimum found in 10a: *m_t_* ∈ {0.05, 0.1, 0.2, 0.4}, the seven values of *D_t_* ≥ 0.03, the three *D_d_*, the eight graphs, IC-B and *r*_3_ = −0.3. The runs with *m_d_* = 0 were taken from 10a. Since *m_d_* lowers the pest load on its own, each efficiency was computed against the no-release run with the same *m_d_*, and the reduction due to *m_d_* was recorded separately.

### Experiment 11, edge-specific transit mortality

This experiment was conditional on Experiment 10 producing a real maximum for at least two values of *m_t_ >* 0 and a relation between D*_t_*\* and *m_t_* that held across graphs. We placed eight nodes uniformly at random in the unit square and joined pairs closer than 0.5 (random geometric graphs), discarding disconnected graphs, until ten connected graphs were collected (seeds 2, 3, 5–9 and 11–13). Transit mortality increased with the Euclidean distance *d_ij_* between patches, *m_ij_* = 1 − exp(−*d_ij_/ℓ*), with *ℓ* ∈ {0.5, 1, 2}, which gave mean edge mortalities of 0.47, 0.28 and 0.15. *D_t_* took 15 log-spaced values between 0.03 and 3, with *D_d_* = 0.03, IC-B and *r*_3_ = −0.3. We compared the observed D*_t_*\* with the value predicted by the relation of Experiment 10, D*_t_*^* =^ *c* |*r*_3_|/(*̄mk_s_*), where *c* is the median of *ρ_s_* over the real maxima of Experiment 10a under IC-B with *r*_3_ = −0.3. for *̄m* we used the mean of *m_ij_* over the edges of the outbreak patch, and alternatively the mean over all edges weighted by the number of parasitoids crossing each edge in the targeted run at D*_t_*\*, Σ*_ij_ A_ij_m_ij_*∫*x*_3*,i*_ d*t/*Σ*_ij_ A_ij_*∫*x*_3*,i*_ d*t*. Weighting only the edges of the outbreak patch would return the simple mean, because its emigrants are split equally among its edges. Finally, we used the mean-degree form 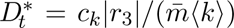, with *̄m* the mean over all edges and *c_k_* the corresponding median of *ρ_k_*.

### (f) Single-patch dynamics and linear stability of the synchronous state

In Experiment 5, to characterise how the release schedule interacts with the intrinsic dynamics, we simulated a single isolated patch, started at (50, 10, 15, 30), for 120 release periods on an 8 × 8 grid with *δ* ∈ {5, 10*, . . .,* 40} and *T* ∈ {5, 8, 11, 15, 20, 25, 30, 40}. The state was sampled immediately after each of the last 30 releases, i.e. on the stroboscopic map of the forced system, and the dispersion of the schedule was defined as the mean Euclidean distance of these 30 four-dimensional states from their centroid. A dispersion near zero indicates a period-*T* response. A small threshold on the dispersion cannot separate a period-*kT* cycle with *k >* 1, whose stroboscopic states are far apart, from an aperiodic response, so we treat the dispersion as a continuous measure of how far the patch is from locking to the release period. We also integrated the patch without releases to *t* = 3000 and estimated the characteristic period of its oscillation from the intervals between successive upward crossings of the mean of *x*_1_ over *t* ∈ [2000, 3000].

Experiment 4 analysed the linear stability of the synchronous state. When releases are uniform and all patches share the same parameters, the synchronous motion **x***_i_*(*t*) = **s**(*t*) for all *i*, where **s** solves the single-patch impulsive system, is an exact solution of equations (2). Its transverse stability can be studied with the master stability function [14, 27]. Projecting the variational equation onto the eigenvectors of *L*, with eigenvalues 0 = *µ*_1_ *< µ*_2_ ≤ · · · ≤ *µ_n_*, decouples it into *n* four-dimensional problems

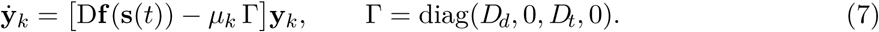

The jump map (3) is a translation, its Jacobian is the identity and it does not enter equation (7). Mode *k* = 1 describes perturbations along the synchronous manifold and modes *k* ≥ 2 describe transverse perturbations.

Because the single-patch trajectory did not settle on a period-*T* orbit (section 3(a)), we estimated the largest Lyapunov exponent of each mode along a long reference trajectory rather than computing Floquet multipliers. The reference trajectory started from (50, 10, 15, 30) with *δ* = 35, *T* = 15 and was integrated to *t* = 30 000. After discarding the first 30% of the run, a unit perturbation vector initially aligned with *x*_1_ was propagated with equation (7) over 1400 consecutive segments of length *T* and renormalised at the end of each segment [28]. The exponent is the mean logarithmic growth rate over all segments, and its standard error was computed from 20 batch means of consecutive segments. A long run was necessary. Over a run to *t* = 1200, with 56 segments, the standard errors were 6–8 × 10*^−^*^3^, larger than most of the exponents, and the estimates were probably still influenced by the slow growth of the oscillation. Estimates from runs to *t* = 12 000 and *t* = 30 000 differed by less than one standard error of the shorter run. We evaluated all transverse modes of the seven seed-1 topologies at *D_d_* = *D_t_* = 0.03 and recorded the largest exponent Λ_max_. As a check, the mode-1 exponent computed this way, with *D_d_* = *D_t_* = 0 and with *D_d_* = *D_t_* = 0.03, was identical in double precision to an independently coded single-patch calculation, and the script used for the long runs reproduced, with a run to *t* = 1200, the exponents of the original stability code to the reported precision. We then compared Λ_max_ with the synchrony index and the spatial coefficient of variation obtained in Experiment 2 on the same seed-1 graphs under the uniform strategy.

With transit mortality the synchronous state exists only on regular graphs, where every node has the same degree *k*. On such a graph *L* = *kI* − *A*, so for an eigenvector **v** of *L* with eigenvalue *µ*, (*L* + *mA*)**v** = (*mk* + (1 − *m*)*µ*)**v**. The synchronous mode (*µ* = 0) then follows the single-patch model with 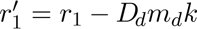 and 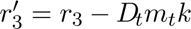, and each transverse mode obeys equation (7) for that modified patch with Γ replaced by diag (1 − *m_d_*)*D_d_,* 0, (1 − *m_t_*)*D_t_,* 0 . Among our graphs only the cycle and the complete graph are regular. For these two we recomputed Λ_max_ with *m_t_* ∈ {0, 0.05, 0.1, 0.2, 0.4}, *m_d_* = 0 and *D_d_* = *D_t_* = 0.03. The reduction was verified by simulating both graphs from a synchronous initial state with *m_d_* = *m_t_ >* 0 and comparing with the modified single patch, which agreed to a relative error below 4 × 10*^−^*^13^.

## 3. Results

### (a) Dynamics of an isolated patch and sensitivity to local parameters

Without releases the single-patch model does not approach an equilibrium. Between *t* = 2000 and *t* = 3000 the psyllid density oscillated between 12.8 and 62.0, with a characteristic period of 7.84 a.u. estimated from 127 successive upward crossings of the mean (s.d. 0.23). The amplitude was still growing during the first 150 a.u. (electronic supplementary material, figure S1*a*), which is the scale of the network experiments.

Periodic releases did not entrain this oscillation. None of the 64 schedules of the single-patch map produced a period-*T* response, and the dispersion of the post-release states ranged from 3.2 to 31.6 (electronic supplementary material, figure S1*b*). Dispersion grew with the release period, from a mean of 5.7 at *T* = 5 to 30.3 at *T* = 40. The lowest value occurred at *δ* = 30, *T* = 15, close to twice the intrinsic period, but the neighbouring schedule *δ* = 35, *T* = 15 used as reference in the network experiments had a dispersion of 28.3. The response to the schedule is therefore sensitive to small changes in release size near this resonance. The largest Lyapunov exponent along the synchronous trajectory with the reference schedule was (0.2 ± 0.4) × 10*^−^*^3^, indistinguishable from zero. This is consistent with quasiperiodic motion, although weak chaos cannot be excluded. In either case, the patch dynamics coupled by the network oscillate persistently.

With the reference configuration of the sensitivity analysis the network AUC was 18 796. Five parameters changed it by more than 9% when perturbed by 10% (electronic supplementary material, figure S2). These were the intrinsic rate of plant vigour *r*_4_ (14.1%), the effect of flush on vigour *b*_42_ (13.7%), the effect of flush on the psyllid *b*_12_ (12.4%), the intrinsic rate of the parasitoid *r*_3_ (12.3%) and the benefit of parasitism to the parasitoid *b*_31_ (9.3%). For the first four the relative change in AUC exceeded the relative change in the parameter, so the output amplifies uncertainty in these inputs. Reducing *r*_4_ by 10% lowered the AUC by 14.1%, and making *r*_3_ 10% more negative, that is increasing parasitoid mortality, raised it by 12.3%. None of the eight crowding coefficients *a_i_* and *c_i_* changed the AUC by more than 1.5%. The responses to *r*_2_ and *b*_21_ were small and had the same sign for both perturbations, which indicates that the AUC is not monotone in these parameters near the reference point.

### (b) Release strategy, size and period

Across the 112 combinations of topology and dispersal rates in Experiment 2, the network AUC without releases ranged from 26 914 to 27 416. Uniform releases lowered it by 30.4–31.0% and releases targeted at the outbreak patch by 2.5–4.5%. The mean psyllid density per patch in the second half of the horizon was 34.0 without releases, 21.9 under uniform releases and 32.6 under targeted releases. These differences mostly reflect the amount of parasitoid released, 1680 units in a uniform programme and 210 in a targeted one.

When efficiency was measured per parasitoid released, the two strategies performed similarly, and which one was better depended on the dispersal rates (figure 2a,b). The AUC avoided per unit released varied little under uniform releases, between 4.87 and 5.05 over all runs. Under targeted releases it varied between 3.25 and 5.79. The targeted strategy was least efficient when psyllids dispersed and parasitoids did not. At *D_t_* = 0 its efficiency, averaged over topologies, fell from 5.30 at *D_d_* = 0 to 3.51 at *D_d_* = 0.1. At *D_t_* = 0.1 it stayed between 5.23 and 5.33 whatever the value of *D_d_*, slightly above the uniform strategy. Targeted releases also left the network more heterogeneous. The mean spatial coefficient of variation of psyllid density was 0.083 under targeted releases against 0.008 under uniform releases and 0.006 without releases.

**Figure 2:**
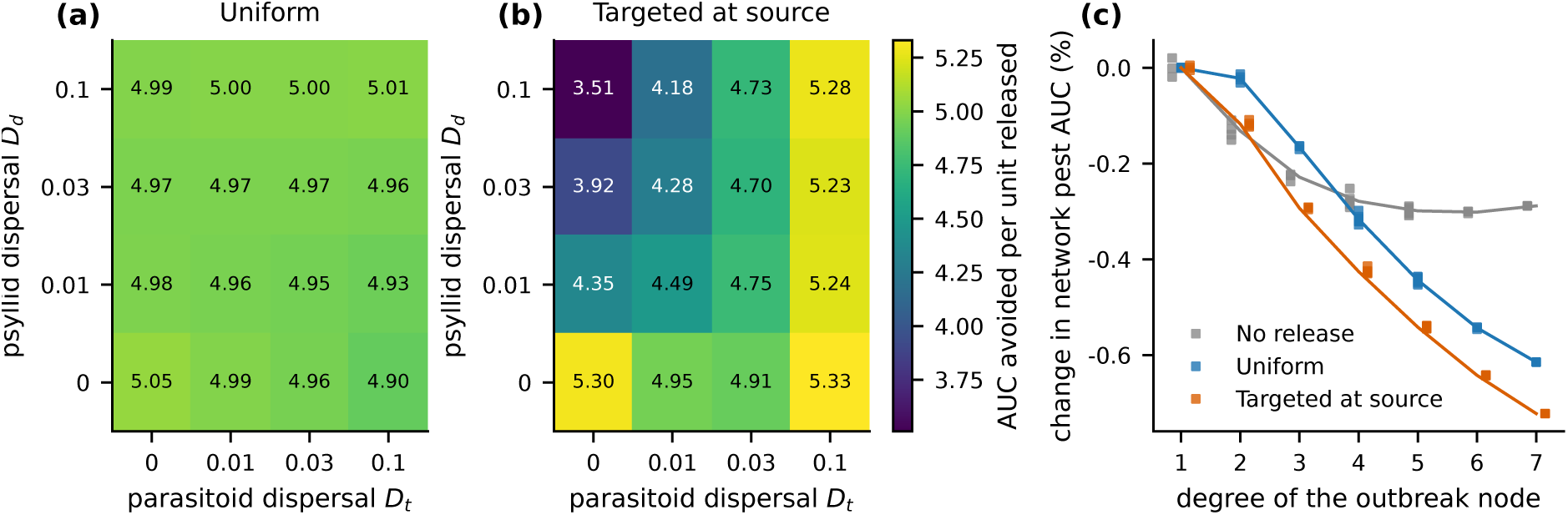
Release strategy and network structure. (*a*,*b*) Network pest load avoided per parasitoid released under uniform (*a*) and targeted (*b*) releases, as a function of the dispersal rates of psyllids (*D_d_*) and parasitoids (*D_t_*). Each cell is the mean over the seven seed-1 topologies of Experiment 2 (*δ* = 35, *T* = 15). (*c*) Change in network pest AUC relative to the mean for graphs whose outbreak patch has degree 1, plotted against the degree of the outbreak patch, for the 34 graphs of Experiment 3 at *D_d_* = *D_t_*= 0.03. Points are individual graphs, slightly offset horizontally by strategy, and lines join the means for each degree.

Dispersal rates had small effects on the pest load. Without releases, raising *D_d_* from 0 to 0.1 at *D_t_* = 0 increased the mean AUC from 27 320 to 27 391 (0.3%), and raising *D_t_*from 0 to 0.1 at *D_d_* = 0 decreased it to 26 983 (1.2%).

Release size and frequency determined how much of the pest load was removed (figure 3a). On the complete graph, uniform releases reduced the AUC by 4.2% with the sparsest and smallest schedule (*δ* = 10, *T* = 30) and by 89.2% with the densest and largest (*δ* = 50, *T* = 5). The path and the random graph reached 89.6% and 89.5% with the latter schedule. Under the most intense schedules psyllid density in some patch fell as low as 0.0013, although it never reached zero within the horizon. Topology changed the reduction achieved by a given schedule by at most 0.4 percentage points under uniform releases and 3.7 points under targeted releases, whose maximum reduction was 14.3% on the star and complete graphs, 11.0% on the random graph and 10.6% on the path.

**Figure 3:**
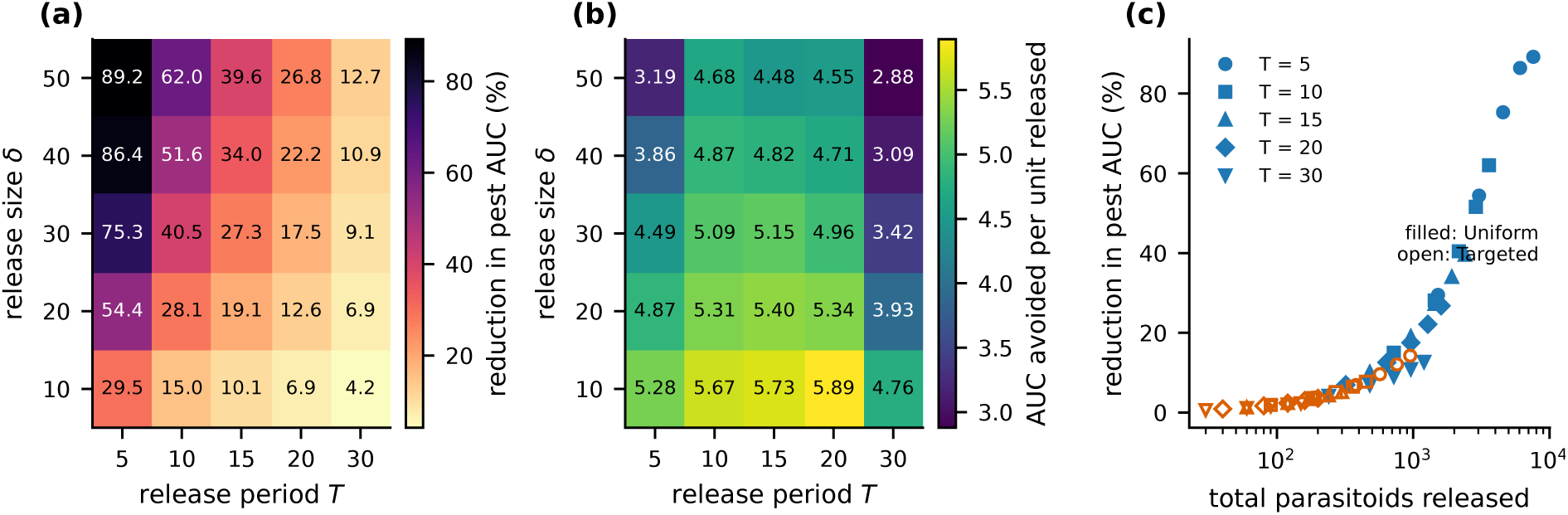
Release size and period (Experiment 6, *D_d_* = *D_t_*= 0.03). (*a*) Reduction in network pest AUC relative to no release and (*b*) pest AUC avoided per parasitoid released, for uniform releases on the complete graph. Efficiencies use the number of releases actually applied, *N* (*T*) = ⌈100*/T* ⌉ − 1. (*c*) Reduction against the total number of parasitoids released on the complete graph, for uniform (filled) and targeted (open) releases. Marker shape indicates the release period.

Efficiency behaved very differently from suppression (figure 3b). For every period it decreased as release size increased, so each additional parasitoid removed less pest load than the previous one. Across all topologies, efficiency ranged from 2.85 to 5.89 under uniform releases and from 3.04 to 6.45 under targeted releases. The most efficient schedule was *δ* = 10, *T* = 20 for all eight combinations of topology and strategy. It achieved reductions of only 6.8–6.9% under uniform releases and 0.9% under targeted releases. This schedule lies at the lower edge of the grid of release sizes, so the maximum of efficiency may well lie outside the range explored and has no particular meaning as an operating point. The longest period tested, *T* = 30, was the least efficient for every release size under uniform releases, with values between 2.85 and 4.76.

When results are plotted against the total number of parasitoids released, the two strategies and all periods fall close to a single increasing, saturating curve (figure 3c). Frequency still mattered at a fixed release rate. Among uniform schedules with *δ/T* = 2 on the complete graph, reductions were 29.5%, 28.1%, 27.3% and 22.2% for *T* = 5, 10, 15 and 20. Part of this decline follows from the finite horizon, since the first release takes place at *t* = *T* and the number of releases *N* (*T*) falls faster than 1*/T*, giving totals of 1520, 1440, 1440 and 1280 parasitoids for the four schedules.

### (c) Network topology and spatial coherence

Network topology had a much weaker effect than the release strategy. For a given pair of dispersal rates, the AUC of the seven seed-1 topologies differed by at most 0.87% without releases, 0.64% under uniform releases and 1.59% under targeted releases. Redrawing the stochastic graphs changed little. Across the ten instances of each stochastic family the coefficient of variation of the AUC never exceeded 0.47%.

The star and the complete graph gave the same results in every run of Experiments 2, 3 and 6, with relative differences below 2 × 10*^−^*^15^, despite algebraic connectivities of 1 and 8. This follows from their structure. When the outbreak patch is adjacent to every other patch and those patches start from the same state, each of them receives the same flow from the outbreak patch and the same releases, so they remain identical at all times. The diffusive flow between two identical patches is zero, so the edges that join them have no effect, and both graphs reduce to an outbreak patch coupled to *n* − 1 identical patches. The argument requires dispersal to conserve individuals. With transit mortality each patch loses individuals at a rate proportional to its own degree, and the equivalence breaks down (section 3(f)).

One Barabási–Albert instance in which node 1 had degree 7 reproduced the star result exactly, as expected. This suggested looking at the degree of the outbreak patch more generally. Across the 34 graphs this single number accounted for nearly all the variation among topologies (figure 2c). Its Spearman correlation with the AUC was −0.98 under both release strategies and −0.87 without releases. Algebraic connectivity and mean degree were weaker predictors, with correlations between −0.33 and −0.42. Moving the outbreak from a patch of degree 1 to one of degree 7 reduced the AUC by 0.61% under uniform releases and by 0.72% under targeted releases. Without releases the relationship was not monotone and the lowest AUC occurred at degree 6. The source degree was also strongly correlated with the synchrony index without releases and under uniform releases (*ρ* = 0.98 in both cases). These effects are consistent and nearly deterministic, but they are small, and in this model the choice of release strategy outweighs topology by more than an order of magnitude. They are also specific to this initial condition and to dispersal rates near 0.03, and they do not survive transit mortality (section 3(f)).

Under uniform releases the largest transverse Lyapunov exponent was negative on all seven topologies (table 3), so the synchronous state is linearly stable on every graph. The path was the least stable, with an exponent about two standard errors below zero. All exponents were small in magnitude. The corresponding e-folding times ranged from 60 a.u. for the complete graph to about 1050 a.u. for the path, against a simulated horizon of 100 a.u. Over that horizon a transverse perturbation would be multiplied by factors between 0.19 and 0.91, so the linear analysis predicts slow convergence everywhere rather than a clear separation between networks. The ranking of the exponents broadly agreed with the ranking of the simulated coherence, with exceptions such as the scale-free graph, more stable than the star but less synchronous. The Spearman correlation between the largest transverse exponent and the synchrony index was −0.86 (*p* = 0.01, *n* = 7), and drawing the exponents from their sampling distributions gave a 95% range of −0.95 to −0.34. Algebraic connectivity was less closely associated with the synchrony index (*ρ* = 0.69, *p* = 0.09). The level of coherence, however, did not follow the size of the exponents. The star and the complete graph had identical synchrony, as expected from the argument above, while their exponents differed by a factor of about seven, and the six graphs other than the complete graph had exponents within a factor of three of each other but synchrony indices between 0.73 and 0.85. Over the horizon relevant to an outbreak, linear stability orders the networks, but the coherence actually reached is set mainly by the initial asymmetry and by how directly the outbreak patch is connected to the rest of the network.

**Table 3:** Largest transverse Lyapunov exponent Λ_max_ of the synchronous state under uniform releases (*δ* = 35, *T* = 15, *D_d_*= *D_t_* = 0.03), and simulated spatial coherence on the same seed-1 graphs. Exponents are means ± standard errors over 1400 segments (section 2(f)). Before rounding, the exponents of the random and small-world graphs are −2.185 and −2.193 × 10*^−^*^3^. *e*^100Λmax^ is the linear amplification of a transverse perturbation over the 100 a.u. horizon.

| topology | λ<sub>2</sub> | 10<sup>3</sup>Λ<sub>max</sub> | *e*<sup>100Λ<sub>max</sub></sup> | synchrony index | spatial CV |
|---|---|---|---|---|---|
| path | 0.152 | −0.95 ± 0.45 | 0.91 | 0.731 | 0.0154 |
| cycle | 0.586 | −1.70 ± 0.47 | 0.84 | 0.744 | 0.0127 |
| star | 1.000 | −2.44 ± 0.47 | 0.78 | 0.851 | 0.0011 |
| complete | 8.000 | −16.61 ± 0.51 | 0.19 | 0.851 | 0.0011 |
| random | 1.074 | −2.19 ± 0.45 | 0.80 | 0.752 | 0.0127 |
| small world | 1.839 | −2.19 ± 0.45 | 0.80 | 0.823 | 0.0015 |
| scale free | 1.067 | −2.61 ± 0.22 | 0.77 | 0.820 | 0.0017 |

### (d) Parasitoid dispersal without transit mortality

Without transit mortality (Experiment 8), the efficiency of targeted releases ranged from 2.98 to 7.09 under IC-A and from 3.02 to 22.82 under IC-B, against 4.68–5.39 and 3.14–8.14 for uniform releases. Its dependence on the dispersal rates differed between the two initial conditions (figure 4a,b). Under IC-A, *E_T_*depended almost only on *D_t_* and rose from 3.3 to 6.0 (median over the seven focal graphs, *r*_3_ = −0.3). Under IC-B it depended on both rates and was highest, up to 10.8, when parasitoids dispersed fast and psyllids slowly. Targeted releases were the more efficient strategy in 56% of the combinations under IC-A and 58% under IC-B. The boundary *E_T_* = *E_U_* lay near *D_t_* ≈ 0.05 under IC-A whatever *D_d_*, and close to the diagonal *D_t_* = *D_d_* under IC-B for *D_d_ <* 0.03. Targeted releases thus paid off once the parasitoid was mobile enough.

**Figure 4:**
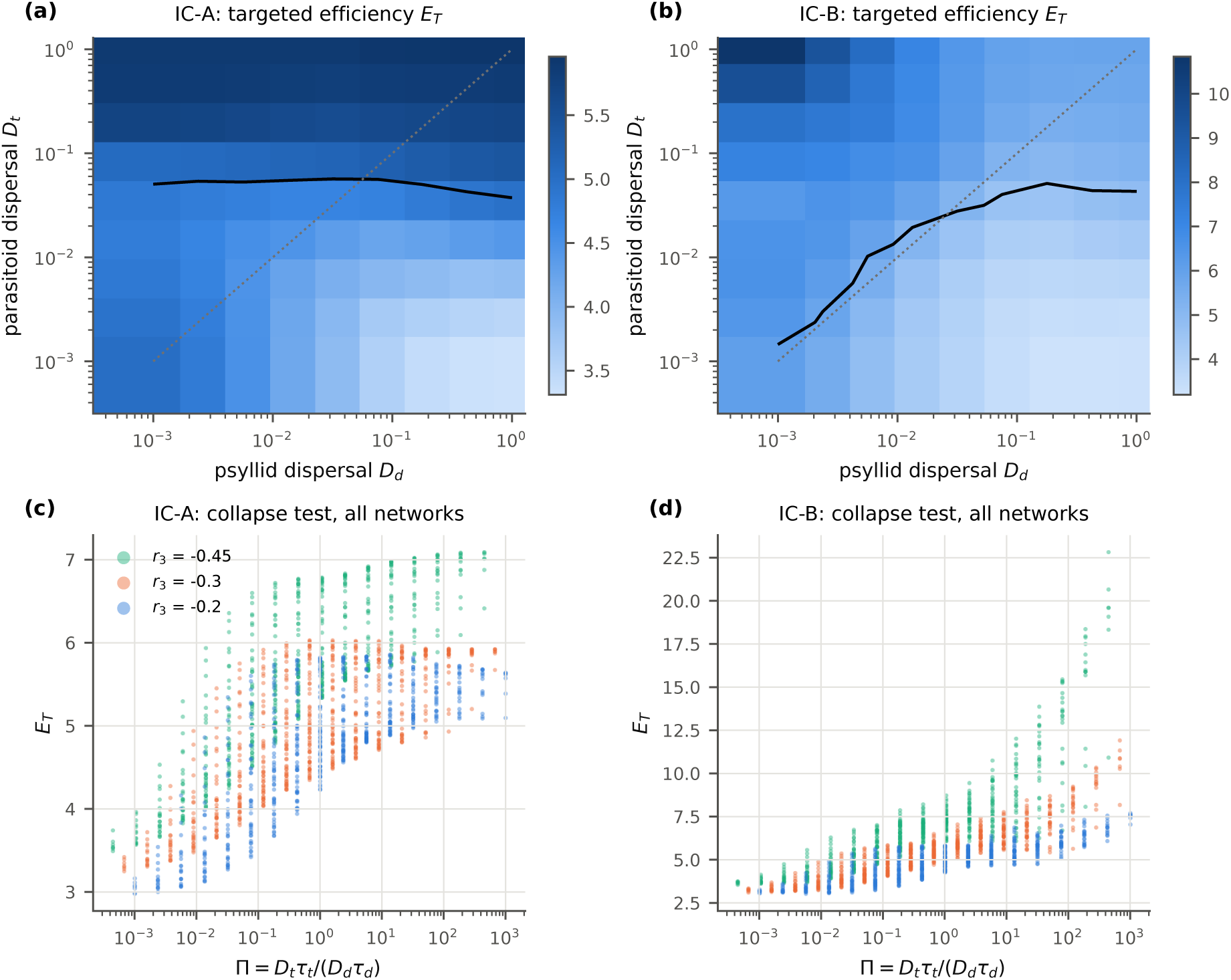
Relative dispersal without transit cost (Experiment 8). (*a*,*b*) Efficiency of targeted releases *E_T_* (pest AUC avoided per parasitoid released) as a function of the dispersal rates of psyllids and parasitoids, under IC-A (*a*) and IC-B (*b*). Each cell is the median over the seven focal graphs with *r*_3_ = −0.3. Black line, *E_T_* = *E_U_* ; dotted line, *D_t_*= *D_d_*. (*c*,*d*) *E_T_* against the relative dispersal Π for all runs, coloured by *r*_3_, under IC-A (*c*) and IC-B (*d*).

No candidate variable collapsed the data (table 4, figure 4c,d). The best cross-validated *R*^2^ was 0.41 under IC-A and 0.18 under IC-B, far from the threshold of 0.9. Even a free function of *D_d_* and *D_t_* reached only 0.52 and 0.18, so most of the variation that a dispersal variable cannot capture is due to *r*_3_, and the rest to the graph. The parasitoid mortality rate did not act only as a time scale. Under IC-B the median *E_T_* was 4.99, 5.56 and 6.71 for *r*_3_ = −0.2, −0.3 and −0.45, because a shorter-lived parasitoid left a much larger pest load to remove, with median AUC_0_ of 18 480, 26 641 and 38 140. Under IC-A the median was 4.89 and 4.86 for the first two values and 5.65 for *r*_3_ = −0.45. Raising |*r*_3_| shortens *τ_t_* and lowers Π, while *E_T_* stays the same or rises, which is why Π performed worse than the simple ratio *D_t_/D_d_*. With *E_T_ /E_U_* as the response the fits improved, to 0.78 for (Π, Π*_t_*) under IC-A, but no candidate reached 0.9. Per-graph maps are given in electronic supplementary material, figure S3.

**Table 4:** Collapse test for the efficiency of targeted releases (Experiment 8, *m_d_* = *m_t_* = 0). *R*^2^ of a cubic fit in log_10_ of each candidate, computed in sample and on held-out data leaving out one value of *r*_3_ or one graph at a time. The two versions of Π differ by a constant factor and give identical fits. The last row of each block is a free bivariate cubic in (log *D_d_,* log *D_t_*), given as an upper reference.

| IC | candidate | in sample | *r*<sub>3</sub> left out | graph left out |
|---|---|---|---|---|
| A | *D*<sub>*t*</sub> | 0.58 | 0.37 | 0.56 |
|  | *D*<sub>*t*</sub>/*D*<sub>*d*</sub> | 0.56 | 0.34 | 0.55 |
|  | Π (*τ*<sub>*d*</sub> = 1/\|*r*<sub>1</sub>\| or *T*<sub>0</sub>) | 0.50 | 0.23 | 0.50 |
|  | (Π, Π<sub>*t*</sub>) | 0.72 | 0.41 | 0.71 |
|  | reference (*D*<sub>*d*</sub>, *D*<sub>*t*</sub>) | 0.74 | 0.52 | 0.71 |
| B | *D*<sub>*t*</sub> | 0.28 | −0.01 | 0.27 |
|  | *D*<sub>*t*</sub>/*D*<sub>*d*</sub> | 0.55 | 0.18 | 0.54 |
|  | Π (*τ*<sub>*d*</sub> = 1/\|*r*<sub>1</sub>\| or *T*<sub>0</sub>) | 0.46 | 0.01 | 0.45 |
|  | (Π, Π<sub>*t*</sub>) | 0.52 | 0.03 | 0.50 |
|  | reference (*D*<sub>*d*</sub>, *D*<sub>*t*</sub>) | 0.58 | 0.18 | 0.55 |

In the absence of transit mortality there was no intermediate parasitoid dispersal rate that maximised the efficiency of targeted releases (Experiment 9, figure 5*a–c*). Under IC-A the maximum of *E_T_* was at the upper end of the grid, *D_t_* = 1, in all 42 curves. Under IC-B it was at *D_t_* = 1 in 33 curves. The other nine, eight of them with *D_d_* = 0.03, had an interior maximum between *D_t_* = 0.15 and 0.85, but with prominence below 2%, and none appeared under both horizons. We therefore treat them as plateaus. *E_T_* was not always monotone, because some curves dipped at low *D_t_*, as in the case *D_d_* = 0 described below, but its largest value was always at, or within 2% of, the fastest dispersal rate tested. Parasitoids that reach patches not yet invaded do not reproduce there, but moving costs them nothing beyond their natural mortality, and they keep dispersing until they find hosts. Uniform releases showed no real maximum either. With *t*_max_ = 300 the curves had small undulations (electronic supplementary material, figure S4), which reflect the phase of the intrinsic oscillation relative to the release schedule rather than numerical error.

**Figure 5:**
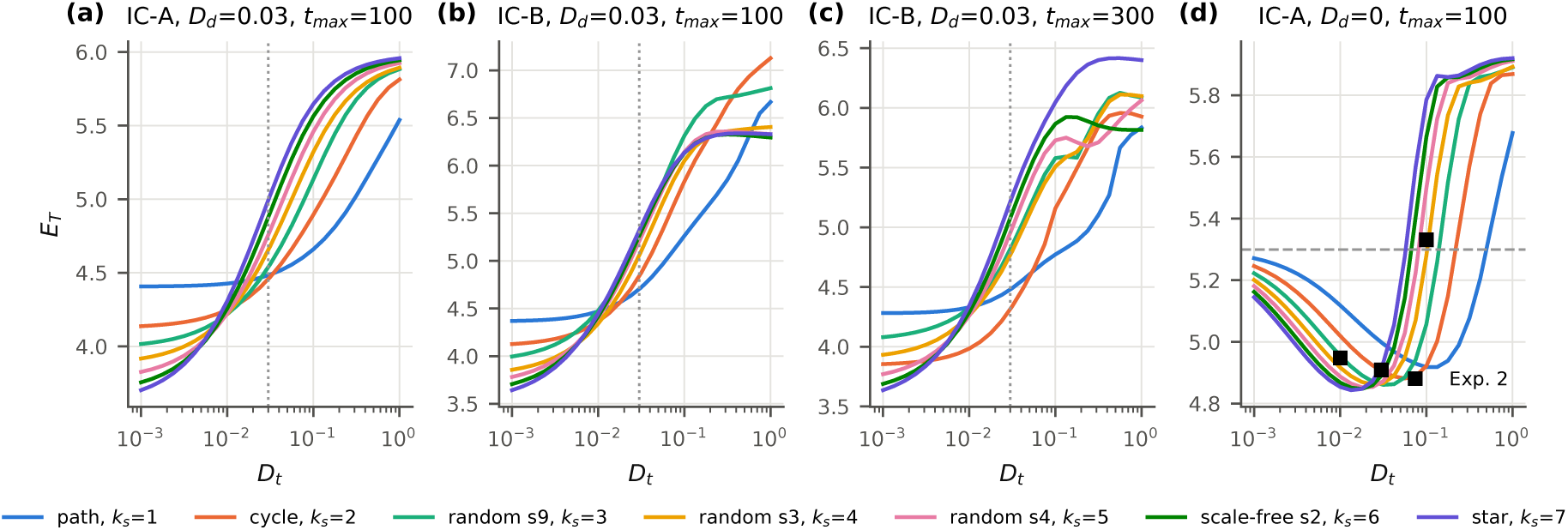
Efficiency of targeted releases against the parasitoid dispersal rate without transit cost (Experiment 9). Lines are the seven focal graphs, labelled by the degree *k_s_* of the outbreak patch. (*a*) IC-A and (*b*) IC-B with *D_d_* = 0.03 and *t*_max_ = 100. (*c*) IC-B with *t*_max_ = 300. Dotted lines mark *D_t_*= *D_d_*. (*d*) Verification run with *D_d_* = 0 and IC-A. The dashed line is *E_T_* at *D_t_* = 0 and squares are the means over the seven topologies of Experiment 2. All conditions are shown in electronic supplementary material, figure S4.

The decrease of *E_T_* in the *D_d_* = 0 row of Experiment 2, from 5.30 at *D_t_* = 0 to 4.91 at *D_t_* = 0.03, was confirmed on a fine grid, where it forms a valley (figure 5*d*). *E_T_* reached a minimum of 4.84–4.92 at *D_t_* between 0.013 (star) and 0.1 (path), and then rose above its value at *D_t_* = 0 from *D_t_* ≈ 0.06 on the star and *D_t_* ≈ 0.5 on the path. The three points of Experiment 2 fall inside the valley. With *t*_max_ = 300 the valley almost disappeared.

### (e) Parasitoid dispersal with transit mortality

When parasitoids died in transit (Experiment 10a), the efficiency of targeted releases developed a pronounced maximum at an intermediate dispersal rate, which moved to lower *D_t_* as *m_t_* increased (figure 6*a–c*; all graphs in electronic supplementary material, figure S5). With *m_t_* = 0 none of the 96 combinations of graph, initial condition, *D_d_* and *r*_3_ had a real maximum. With *m_t_ >* 0, 127 of 384 curves had one, 68 under IC-B and 59 under IC-A, with a median prominence of 73% (range 6–97%). The median efficiency at the maximum was 16.1, against 6.9 at *D_t_* = 1 without mortality. On the complete graph under IC-B (*r*_3_ = −0.3, *D_d_* = 0.03) the optimum was at *D_t_*\* = 0.59, 0.30 and 0.17 for *m_t_*= 0.1, 0.2 and 0.4. On graphs with few edges, such as the path and the star, the optimum lay beyond *D_t_* = 1 in almost every curve with *m_t_* ≤ 0.2 and in about half of those with *m_t_* = 0.4.

**Figure 6:**
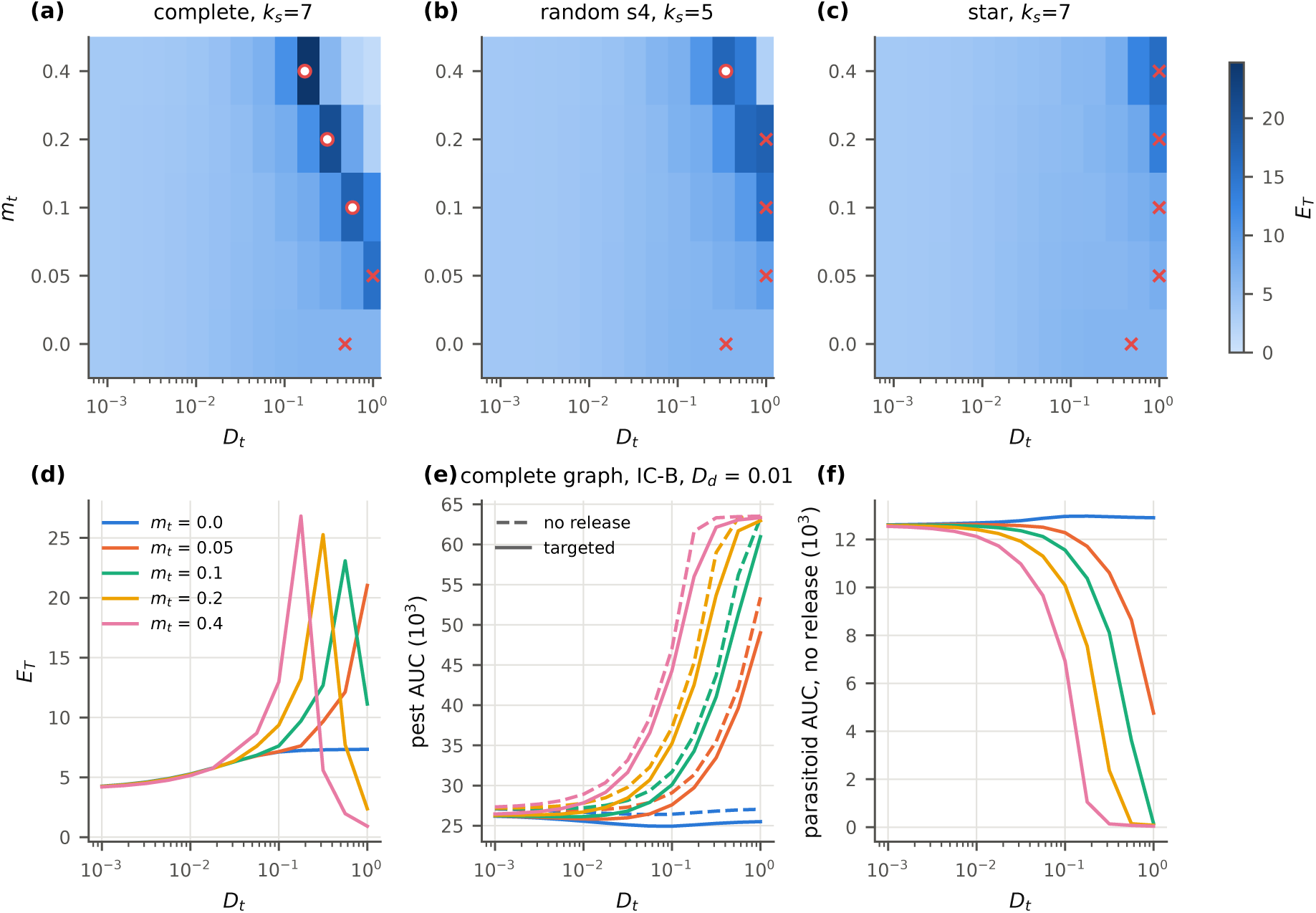
Transit mortality of the parasitoid (Experiment 10a, IC-B, *r*_3_ = −0.3, *m_d_* = 0). (*a*–*c*) Efficiency of targeted releases on the plane (*D_t_*, *m_t_*) for the complete graph, a Gilbert graph with *k_s_* = 5 and the star, with *D_d_* = 0.03. Open circles, interior maxima with prominence above 5%; crosses, maxima at the edge of the grid or with lower prominence. (*d*–*f*) Complete graph with *D_d_* = 0.01. (*d*) *E_T_* . (*e*) Pest AUC with targeted releases (solid) and without releases (dashed). (*f*) Parasitoid AUC without releases. Colours indicate *m_t_*.

The maximum is not a minimum of the pest load, which in most curves was lowest at or near the lowest *D_t_* (figure 6*d–f*). With *m_t_ >* 0 the pest AUC under targeted releases increased with *D_t_*. When it had an interior minimum, which happened in 53% of the curves, the minimum was a median of 0.9% below the lower of the two ends of the curve, but up to 15% below it on the path and 9% on the cycle under IC-B. The optimum arises from the no-release reference. As *D_t_*increases, transit losses drive the resident parasitoid population towards extinction and the pest load without releases rises steeply, from about 27 000 to 63 500 on the complete graph. At *D_t_*\* the parasitoid AUC without releases was 2–40% of its value with *m_t_* = 0 (median 20%). Releases still maintained a parasitoid population there, so each released parasitoid avoided a large pest load. At higher *D_t_* the released parasitoids also died in transit and *E_T_* fell towards zero. The optimum is therefore the dispersal rate at which releases make the largest difference relative to not releasing.

The optimum fell roughly where transit losses at the outbreak patch matched the natural mortality of the parasitoid, but the agreement was only to an order of magnitude (figure 7*a*). Over the 127 real maxima, 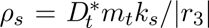 had a median of 1.36 and ranged from 0.49 to 5.66. For a given graph it was nearly constant, and *D_t_*\* decreased almost as 1*/m_t_*, with a median log–log slope of −0.91 (interquartile range −0.95 to −0.87, IC-B). Most of the spread was between graphs. The ratio was about 3.4 on the star, whose hub has degree 7 but whose other patches have degree 1, and about 0.5 on the path. Replacing the source degree by the mean degree of the graph reduced the spread (figure 7*b*). 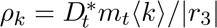 had a median of 0.89 and ranged from 0.60 to 2.05. The standard deviation of log_10_ *ρ* fell from 0.23 to 0.16, which halves its variance. This is consistent with the mechanism above, in which the optimum follows the collapse of the resident parasitoids in the whole network and not only at the outbreak patch.

**Figure 7:**
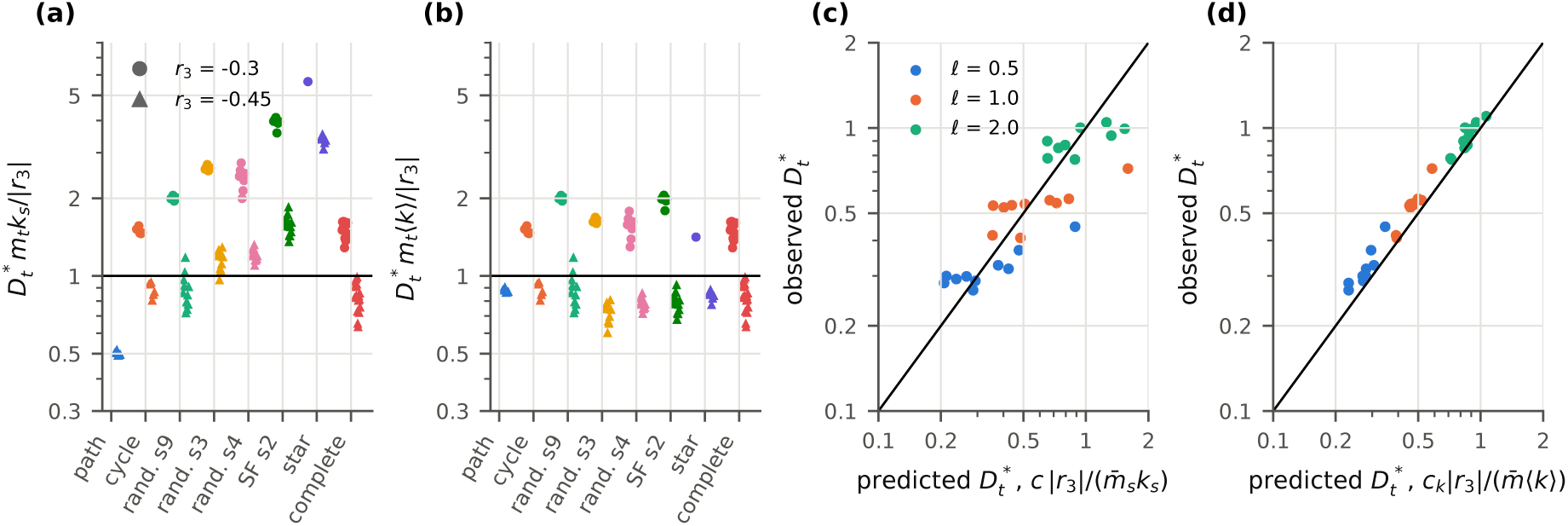
Location of the optimal parasitoid dispersal rate. (*a*) *ρ_s_*= *D^∗^m_t_k_s_/*|*r*_3_| and (*b*) *ρ_k_* = *D^∗^m_t_*⟨*k*⟩*/*|*r*_3_| for the 127 interior maxima with prominence above 5% of Experiment 10a (both initial conditions, three *D_d_*, four *m_t_ >* 0), grouped by graph. Circles, *r*_3_ = −0.3; triangles, *r*_3_ = −0.45. The horizontal line is the ratio of 1 implied by equal transit and natural mortality. (*c*,*d*) Experiment 11. Observed optimum against the prediction from Experiment 10 using the source degree and the mean mortality of the edges of the outbreak patch (*c*), or the mean degree and the mean mortality of all edges (*d*). Colours indicate the characteristic distance *ℓ* of the mortality kernel. The line is the identity.

The dependence on *r*_3_ was opposite to the one implied by the ratio. Under IC-B the transit loss rate at the optimum, 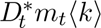, was 0.46 (range 0.39–0.60) with *r*_3_ = −0.3 and 0.35 (0.27–0.41) with *r*_3_ = −0.45. Under IC-A it was 0.49 and 0.39. A parasitoid with higher natural mortality therefore tolerated a lower transit loss before its resident population collapsed. These values are consistent with a persistence threshold for the resident parasitoid, *D_t_m_t_*⟨*k*⟩ ≈ *b*_31_*̄x*_1_ − |*r*_3_|, where *̄x*_1_ is the psyllid density once the parasitoid has collapsed (79.5 per patch under IC-B and 86 under IC-A in our runs). The threshold gives 0.50 and 0.35 under IC-B and 0.56 and 0.41 under IC-A, and section 4 motivates it with a linear invasion argument.

Experiment 10 met the condition for Experiment 11. Under IC-B there was a real maximum for at least two values of *m_t_ >* 0 in 19 of 48 combinations of graph, *D_d_* and *r*_3_, spread over five graphs, and the slope of *D_t_*\* against *m_t_* was similar across them. With distance-dependent mortality on random geometric graphs, all 30 curves had a real maximum, with prominences of 89–98% (figure 7*c,d*). The relation from Experiment 10 with the source degree (*c* = 1.96) and the mean mortality of the edges of the outbreak patch predicted *D^∗^* with a median error of 22% and a correlation of 0.86 on log scales. Weighting all edges by the number of parasitoids crossing them did not improve the prediction (median error 24%, correlation 0.88). The mean-degree form (*c_k_* = 1.54, mean mortality over all edges) had a median error of 10%, a bias of +10% and a correlation of 0.99. Within each value of *ℓ* its correlation with the observed optimum was 0.88–0.97, against 0.76–0.91 for the source-degree form. *D_t_*\* varied by less than a factor of 1.8 within each *ℓ* while *k_s_* ranged from 2 to 6, so the degree of the outbreak patch had little influence on the optimum.

### (f) Network structure with transit mortality

The equivalence between the star and the complete graph no longer held once parasitoids died in transit (figure 8*a*). The difference between the star and the complete graph was zero for *m_t_* = 0. With *m_t_ >* 0 the star always had the lower pest AUC, by up to 36–42% at high *D_t_*depending on *m_t_* and on the initial condition. In the complete graph every patch has degree 7 and loses parasitoids along seven edges, whereas the leaves of the star have a single edge.

**Figure 8:**
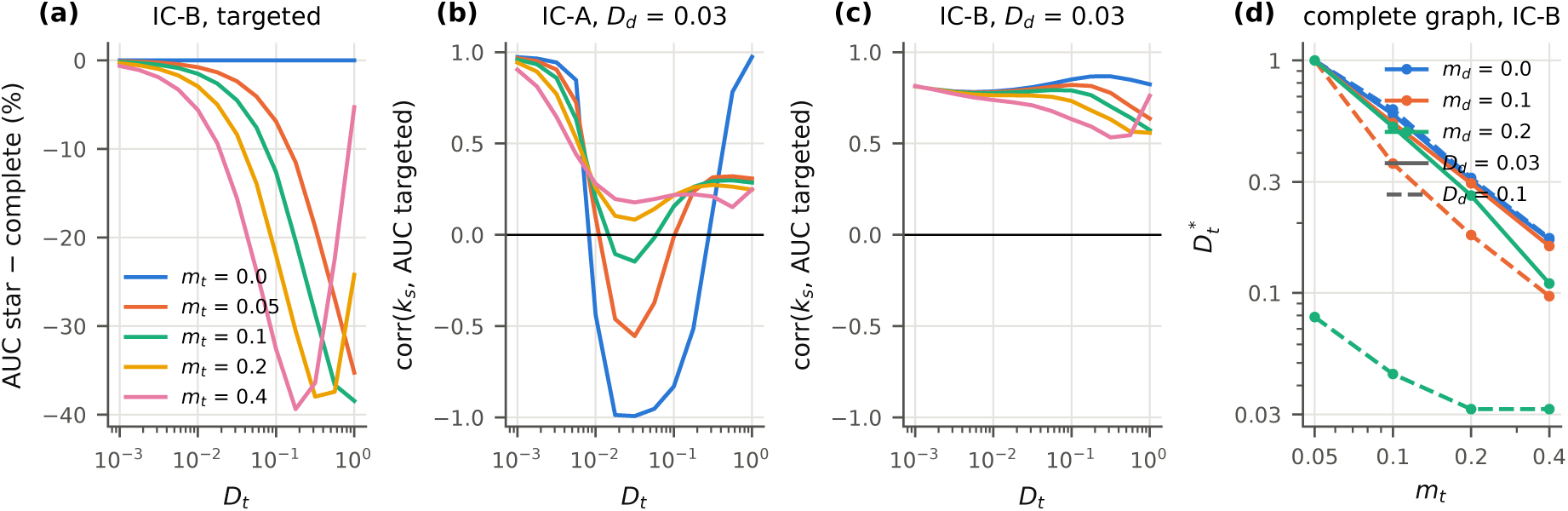
Consequences of transit mortality for the structural results. (*a*) Difference in pest AUC between the star and the complete graph under targeted releases, relative to the complete graph (IC-B, *D_d_*= 0.03, *r*_3_ = −0.3). (*b*,*c*) Pearson correlation across the seven focal graphs between the degree of the outbreak patch and the pest AUC under targeted releases, under IC-A (*b*) and IC-B (*c*), with *D_d_* = 0.03. Colours in (*a*–*c*) indicate *m_t_*. (*d*) Optimal parasitoid dispersal rate on the complete graph against *m_t_* for three values of the psyllid transit mortality *m_d_*, with *D_d_* = 0.03 (solid) and 0.1 (dashed). The optimum was searched for *D_t_* ≥ 0.03 only (Experiment 10b).

The negative association between the degree of the outbreak patch and the pest load also depended on the absence of transit costs (figure 8*b,c*). Across the seven focal graphs under IC-A, with *D_d_* = 0.03 and *D_t_* ≈ 0.03, the Pearson correlation between *k_s_* and the AUC under targeted releases was −0.99 with *m_t_* = 0, in line with section 3(c), and −0.55, −0.15, +0.08 and +0.18 with *m_t_* = 0.05, 0.1, 0.2 and 0.4. Its sign changed between *m_t_* = 0.1 and 0.2. Even without mortality the correlation was strongly negative only for *D_t_* between about 0.02 and 0.1 and became positive for very low or very high *D_t_*. Under IC-B it was positive in every condition (0.40 to 0.88). A well-connected outbreak patch then spreads a pest that was absent elsewhere, and this outweighs its advantage for dispersing parasitoids.

On the two regular graphs, where a synchronous state still exists, transit mortality did not destabilise it. With *D_d_* = *D_t_* = 0.03 the largest transverse exponent stayed negative for every *m_t_*. On the cycle it was −1.70 × 10*^−^*^3^ at *m_t_* = 0 and between −1.27 × 10*^−^*^3^ and −1.80 × 10*^−^*^3^ for *m_t_* = 0.05–0.4. On the complete graph it was −16.6 × 10*^−^*^3^ at *m_t_* = 0 and between −13.0 × 10*^−^*^3^ and −16.3 × 10*^−^*^3^ with mortality. Transit mortality therefore changed the exponents by at most about 25%, without a consistent direction. Standard errors were below 0.33 × 10*^−^*^3^ for every *m_t_ >* 0.

Transit mortality of the psyllid (Experiment 10b) always moved the optimum to lower *D_t_* (figure 8*d*; electronic supplementary material, figure S6). The median shift was −2% and −4% for *m_d_* = 0.1 and 0.2 with *D_d_*= 0.01, −3% and −11% with *D_d_*= 0.03, and −18% and −45% with *D_d_* = 0.1. The shift grew with the psyllid loss rate *D_d_m_d_*. In the most extreme case, the complete graph with *D_d_* = 0.1 and *m_d_* = 0.2, the optimum reached the lower edge of the reduced grid, so its position there is only an upper bound. Without releases, psyllid transit mortality reduced the pest load by a median of 1.8% for *m_d_* = 0.1 (maximum 30%) and 3.6% for *m_d_* = 0.2 (maximum 60%).

## 4. Discussion

Extending the single-patch model of Rodríguez-Falcón et al. [5] to a network of patches changed less than the network literature might lead one to expect, as long as dispersal was free. The amount of parasitoid released, and therefore the release strategy, determined the pest load far more than topology did, and the degree of the outbreak patch, although a nearly deterministic predictor, moved the network AUC by less than one per cent. Once parasitoids died while moving, the efficiency of targeted releases acquired a pronounced maximum at an intermediate dispersal rate, its position scaled with the transit losses summed over the whole network, and two of the structural results obtained without mortality, the equivalence of the star and the complete graph and the benefit of a well-connected outbreak patch under a background infestation, no longer held.

### Release strategy, network structure and the outbreak patch

Uniform releases lowered the network pest load by about 30% and targeted releases by 2.5–4.5%, but a uniform programme used eight times as many parasitoids. Per parasitoid released the two strategies were comparable, and the choice between them depended on how fast each species moved. Concentrating releases at the outbreak patch paid off once the parasitoid was mobile enough (section 3(d)). In both cases targeting was clearly worse when psyllids moved and parasitoids did not. In epidemic models the degree of the seed node and the targeting of hubs can decide whether a perturbation invades the network [11, 12], and dispersal network structure can alter metapopulation dynamics strongly even in simple ecological models [29, 30]. In our model the corresponding effects were consistent but small. A likely reason is that every patch carries the same local dynamics and that the dispersal rates of Experiments 2 and 3, at most 0.1, couple the patches weakly over the horizon of an outbreak, so topology can only modulate how fast the initial asymmetry is redistributed. The equivalence of the star and the complete graph shows that, when the outbreak patch is adjacent to all others, the edges among the remaining patches are dynamically irrelevant. This argument covers only that case, but across the 34 graphs of Experiment 3 the source degree also summarised how directly the outbreak was coupled to the rest of the network.

The negative association between source degree and pest load held only with a background infestation (IC-A), dispersal rates near 0.03 and no transit mortality. Under a point introduction (IC-B) a well-connected outbreak patch spread a pest that was absent elsewhere and the association became positive, and with parasitoid transit mortality the sign reversed already at *m_t_* ≈ 0.2.

### Spatial coherence over the horizon of an outbreak

The master stability function [14, 27] predicted that the synchronous state is linearly stable on all seven topologies, and the ranking of the transverse exponents broadly agreed with the ranking of the observed synchrony. Their magnitudes, however, imply e-folding times between 60 and about 1050 time units, against a simulated horizon of 100, so over an outbreak a transverse perturbation shrinks only by a factor between 0.19 and 0.91. The star and the complete graph, whose exponents differ by a factor of about seven, had identical synchrony. The linear analysis therefore orders the networks but does not set the level of coherence, which depended on the initial asymmetry and on the way the outbreak patch was coupled to the rest, a transient property rather than an asymptotic one [31]. On the random and small-world graphs the largest transverse exponent came from an intermediate eigenvalue (*µ* = 3.27 and 3.31) rather than from *µ*_2_ (table 3), so *λ*_2_ alone does not determine transverse stability in this model.

The exponents are also hard to estimate. Over a trajectory of 840 time units their sampling errors were larger than most of the exponents, and the path even appeared unstable, so only a much longer trajectory resolved their signs. The local dynamics add to this, because the isolated patch oscillates with a period close to 7.8 time units and periodic releases did not entrain it. With patch-level oscillations that are locked neither to the release calendar nor, over an outbreak, to each other, a familiar setting for spatial synchrony in natural populations [32], the asymptotic stability of a synchronous orbit is a weak guide to an outbreak of finite duration.

### Transit mortality and an optimal dispersal rate

Heimpel and Asplen [33] proposed that biological control agents should disperse neither too little nor too much. Our results support a restricted version of this idea. Without a cost of dispersal there was no intermediate optimum. Parasitoids that left the outbreak patch were not lost, since moving carried no cost beyond natural mortality, and the efficiency of targeted releases saturated at high dispersal rates instead of declining. With transit mortality, a cost documented for many dispersing organisms [34], a maximum appeared in a third of the curves, with a median prominence of 73%, and moved to lower *D_t_* almost as 1*/m_t_*. Most of the remaining curves peaked at the upper edge of the grid, *D_t_* = 1, so the third is a lower bound set by the range explored.

The maximum is not a minimum of the pest load. With transit mortality the pest load under targeted releases tended to rise with *D_t_*. Its minimum was usually at or near the lowest dispersal rate, although after a point introduction interior minima reached 15% below the lower end of the curve on the path and 9% on the cycle. The optimum arose because the no-release reference deteriorated. As *D_t_* increased, transit losses drove the resident parasitoid population towards extinction and the pest load without releases more than doubled on the complete graph, while repeated releases still sustained parasitoids near the outbreak patch. Each released parasitoid then avoided a large pest load. The optimum identifies the conditions under which releases make the largest difference relative to doing nothing, and those are conditions under which natural control has already failed. An efficiency defined against a counterfactual can therefore be highest in landscapes that are the worst for the parasitoid, and comparisons of release programmes across sites should report the pest load achieved together with the efficiency.

The scaling of the optimum with the mean degree ⟨*k*⟩, rather than with the degree of the outbreak patch, and its unexpected dependence on *r*_3_ can both be understood from a linear invasion argument. Near *x*_3_ = 0 the crowding term of equation (1) vanishes, and if the psyllid density *x*_1_(*t*) is the same in every patch, the parasitoid vector obeys **ẋ** _3_ = [*r*_3_ + *b*_31_*x*_1_(*t*)] **x**_3_ − *D_t_* (*K* − (1 − *m_t_*)*A*) **x**_3_. The first term is a scalar multiple of the identity and commutes with the second, so a resident parasitoid population at low density grows or declines at the rate

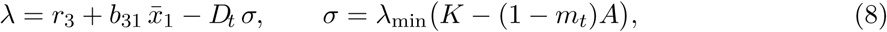

where *̄x*_1_ is the time average of the psyllid density in the absence of the parasitoid. Because the uniform vector is the null vector of *L*, first-order perturbation theory gives 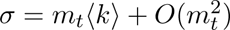, and the Rayleigh quotient gives *σ* ≤ *m_t_*⟨*k*⟩ exactly, with equality on regular graphs. For the eight graphs of Experiment 10, *σ/*(*m_t_*⟨*k*⟩) lies between 0.91 and 1 for *m_t_* ≤ 0.1 and between 0.65 and 1 for *m_t_* = 0.4. Setting *λ* = 0 yields the threshold *D_t_m_t_*⟨*k*⟩ ≈ *b*_31_*̄x*_1_ − |*r*_3_| suggested empirically in section 3(e), which gives 0.50 and 0.35 for *r*_3_ = −0.3 and −0.45 under IC-B, against 0.46 and 0.35 observed, and 0.56 and 0.41 under IC-A, against 0.49 and 0.39. Because *σ* ≤ *m_t_*⟨*k*⟩, equation (8) places the collapse of the resident parasitoid at or above these values. The optima nevertheless lay slightly below them, where the resident population was already declining. The likely reason is that *̄x*_1_, measured once the parasitoid has gone, overestimates the host density available to the parasitoid while it is still present and suppressing the psyllid, which lowers the effective threshold. Equation (8) also explains why a larger |*r*_3_| lowers the tolerable transit loss, since the parasitoid has less margin for growth before dispersal costs are added. With edge-specific mortalities the same first-order argument gives *σ* ≈ ⟨*k*⟩ *̄m*, with *̄m* the mean of *m_ij_* over all edges, which is the mean-degree form that predicted the optimum of Experiment 11 with a median error of 10%. Psyllid transit mortality lowers *̄x*_1_, and with it the threshold, which is consistent with the shift of *D_t_*\* to lower values in Experiment 10b. The argument is a heuristic. It assumes a spatially uniform psyllid density, which IC-B violates early in the outbreak, and it locates the collapse of the resident population rather than the efficiency maximum itself, which also depends on the finite horizon. It predicts, however, that on irregular graphs *ρ_k_* should drift upwards at large *m_t_*, where *σ* falls below *m_t_*⟨*k*⟩. The stored output does not support this refinement. *ρ_k_* rose with *m_t_* on the cycle and the complete graph as well, where *σ* = *m_t_*⟨*k*⟩ exactly, and replacing *m_t_*⟨*k*⟩ by *σ* did not reduce the spread of the ratio (standard deviation of its log_10_, 0.159 against 0.155). The mean-degree form is therefore as good a summary as the exact eigenvalue, and the residual drift with *m_t_* has another origin, possibly the finite horizon. The quantity *σ* plays the same role as the leading eigenvalue that defines the metapopulation capacity of a landscape [35], a landscape-level property that combines topology and movement costs into a single persistence threshold.

The failure of every dimensionless group to collapse the efficiencies in Experiment 8 points in the same direction. Most of the variance that dispersal rates could not explain was due to *r*_3_, which changes how much pest there is to remove and not only how long a parasitoid lives. A predictive scaling variable would have to include a measure of the pest load available to be suppressed, such as AUC_0_ or *b*_31_*̄x*_1_*/*|*r*_3_|, alongside the dispersal time scales.

### Implications for release programmes against *Diaphorina citri*

Augmentative biological control is often judged on cost per unit of pest suppressed [36, 37], and on that criterion our results show that efficiency fell with release size and that targeting the outbreak patch was competitive only above a threshold of parasitoid mobility. The most efficient schedule lay at the edge of the grid and achieved reductions below 7%, so the efficiency optimum is not an operating point, and programmes have to trade suppression for efficiency explicitly. After a point introduction, the mobility threshold for targeting rose with the mobility of the psyllid. *D. citri* can move between groves and over long distances [38, 39], whereas the dispersal radius of released *T. radiata* measured in Brazilian groves was about 18 m [40]. Model rates are in arbitrary units and cannot be matched directly to these measurements, but if released parasitoids stay within about 20 m of the release point while psyllids move between groves, targeted releases into a single focus are likely to lose to area-wide releases, above all after a point introduction, which agrees with the emphasis on coordinated regional management of HLB [41].

Release accuracy and host-dependent searching also matter, as other spatially explicit release studies have stressed [6, 7, 10]. *Telenomus podisi* left host-rich plants more slowly, and releases placed on infestation foci raised parasitism [9]. Our parasitoids dispersed at a constant rate irrespective of host density. Host-dependent emigration would retain parasitoids in infested patches and reduce transit losses, which would probably move the optimum to higher *D_t_* for real parasitoids with directed search.

### Limitations

The model is formulated in arbitrary units and its parameters were not estimated from field data, so our conclusions are qualitative. The networks had eight patches and dispersal was linear and density-independent. Larger landscapes, distance-dependent rates and host-dependent movement [8, 42] could further change the role of topology. The model is deterministic, so psyllid densities that fell to 10*^−^*^3^ under intense schedules would correspond to local extinction in a stochastic model, and our results say nothing about eradication. The sensitivity analysis was local and one-at-a-time and explored only a neighbourhood of the reference point [26]. Several conclusions depended on the initial condition, and the horizon of 100 time units, chosen to represent an outbreak, is short relative to most transverse time scales and to the growth of the intrinsic oscillation. Finally, the crowding terms of the local model can become negative, so boundedness relies on interaction terms, a feature inherited from the single-patch formulation [5] that limits the range of release schedules over which the model can be trusted.

## 5. Conclusions

Moving from one patch to a network did not, by itself, change how augmentative releases of *T. radiata* should be judged. When dispersal was free, the number of parasitoids released set the pest load, uniform and targeted releases were comparable per parasitoid, and the graph was a minor modifier whose effect ran mainly through the degree of the outbreak patch. The linear stability of the synchronous state ordered the networks by their spatial coherence, but the level of coherence reached over an outbreak was set mainly by transient dynamics.

Transit mortality made network structure matter. It produced an optimal parasitoid dispersal rate for targeted releases, it broke the equivalence between the star and the complete graph, and it reversed the benefit of a well-connected outbreak patch seen under a background infestation. The optimum follows a persistence threshold for the resident parasitoid that depends on the mean degree and the mean edge mortality rather than on the outbreak patch alone. Because it marks the point where natural control fails, a high efficiency per released parasitoid can coexist with a high pest load, and programmes should report both.

For *D. citri*, these results favour area-wide releases when parasitoids move little, either in absolute terms or, after a point introduction, relative to the psyllid, and give no support to targeting the best-connected grove as a general rule. Two measurements would decide which regime applies to citrus landscapes, the relative dispersal rates of psyllid and parasitoid between groves and the mortality of *T. radiata* while moving between them. A calibrated, stochastic version of the network model, together with a global sensitivity analysis, would be needed before these conclusions are translated into release schedules. The local analysis already points to plant vigour and flush as the main sources of uncertainty. A spatial optimal-control formulation could then choose the timing, size and location of releases under a fixed budget, rather than comparing prescribed schedules.

## Data accessibility

The Python code and the CSV outputs of every experiment are available at https://github.com/ sebrodfal/DiaphorinaNetworkModel [25]. The repository includes the scripts that regenerate all figures and derived quantities from those outputs.

