## Supplementary Figures S1-S6 for "Transit mortality and the efficiency of *Tamarixia radiata* releases against *Diaphorina citri* in metapopulation networks"

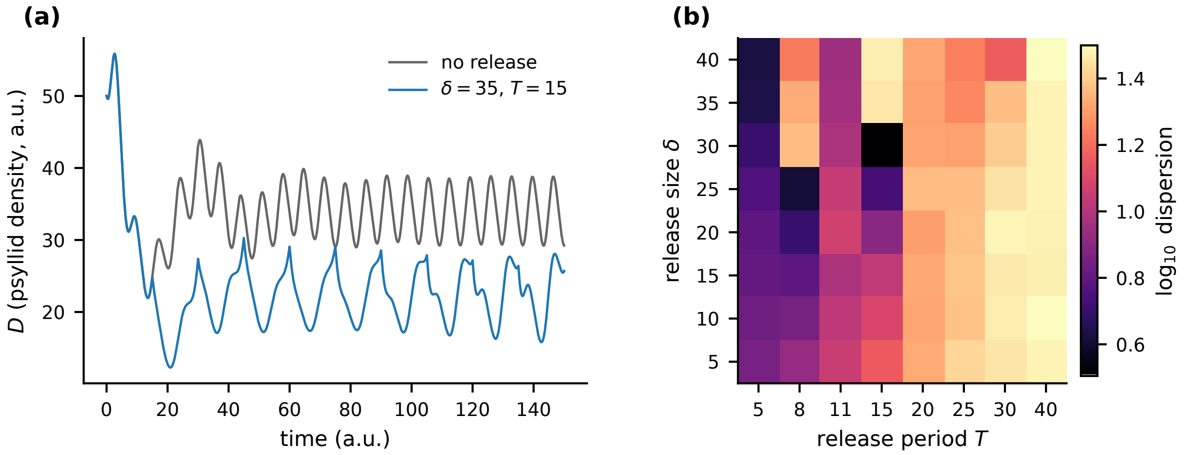


**Figure S1:** Dynamics of an isolated patch. ($a$) Psyllid density without releases (grey) and with releases of $\delta = 35$ every $T = 15$ (blue), from the initial state $(50, 10, 15, 30)$. ($b$) Dispersion of the state sampled after each of the last 30 of 120 releases, on a $\log_{10}$ scale. Zero dispersion would indicate a response of period $T$. No schedule on the grid reached a dispersion below 3.2.

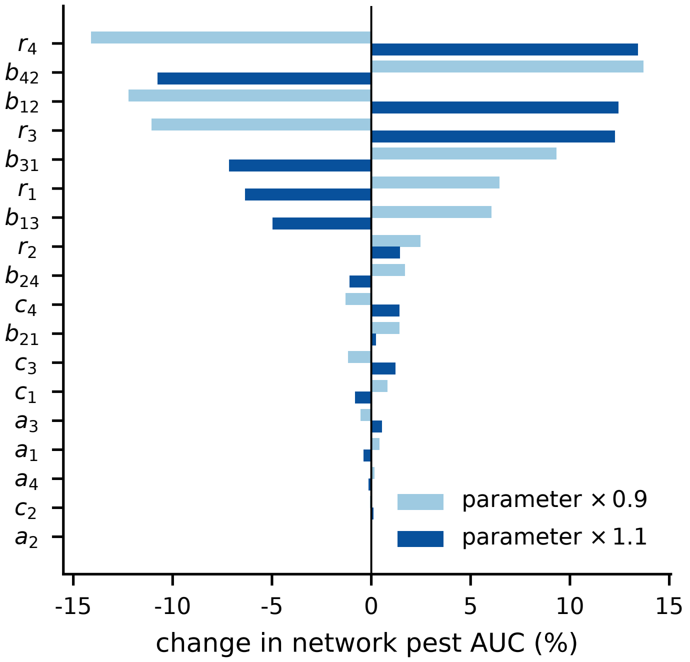


**Figure S2:** One-at-a-time sensitivity of the network pest AUC to multiplying each local parameter by 0.9 or 1.1. Complete graph, uniform releases, $\delta = 35$, $T = 15$, $D_d = D_t = 0.03$. Parameters are ordered by the larger absolute change. For negative parameters the factor 1.1 increases their magnitude.

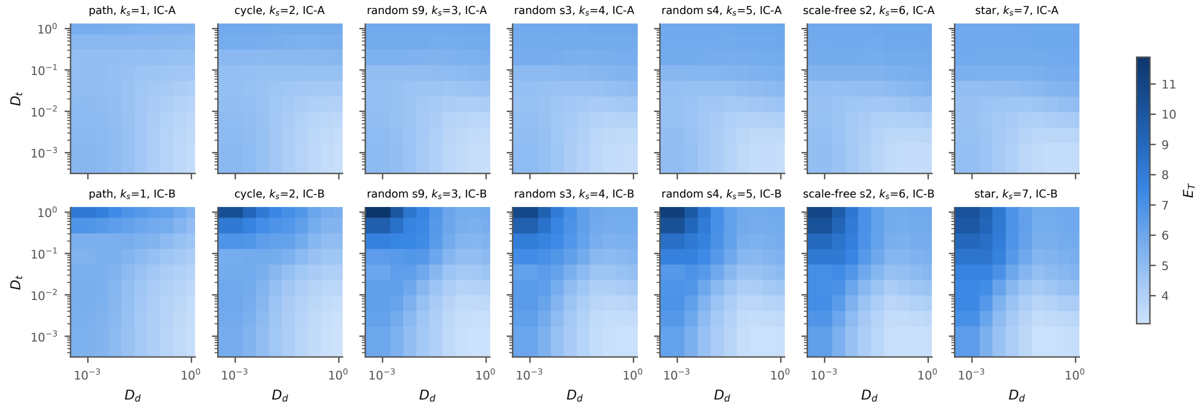


**Figure S3:** Efficiency of targeted releases $E_T$ (pest AUC avoided per parasitoid released) as a function of the dispersal rates of psyllids ($D_d$) and parasitoids ($D_t$) for each of the seven focal graphs, under IC-A (top row) and IC-B (bottom row), with $r_3 = -0.3$ (Experiment 8). All panels share the colour scale.

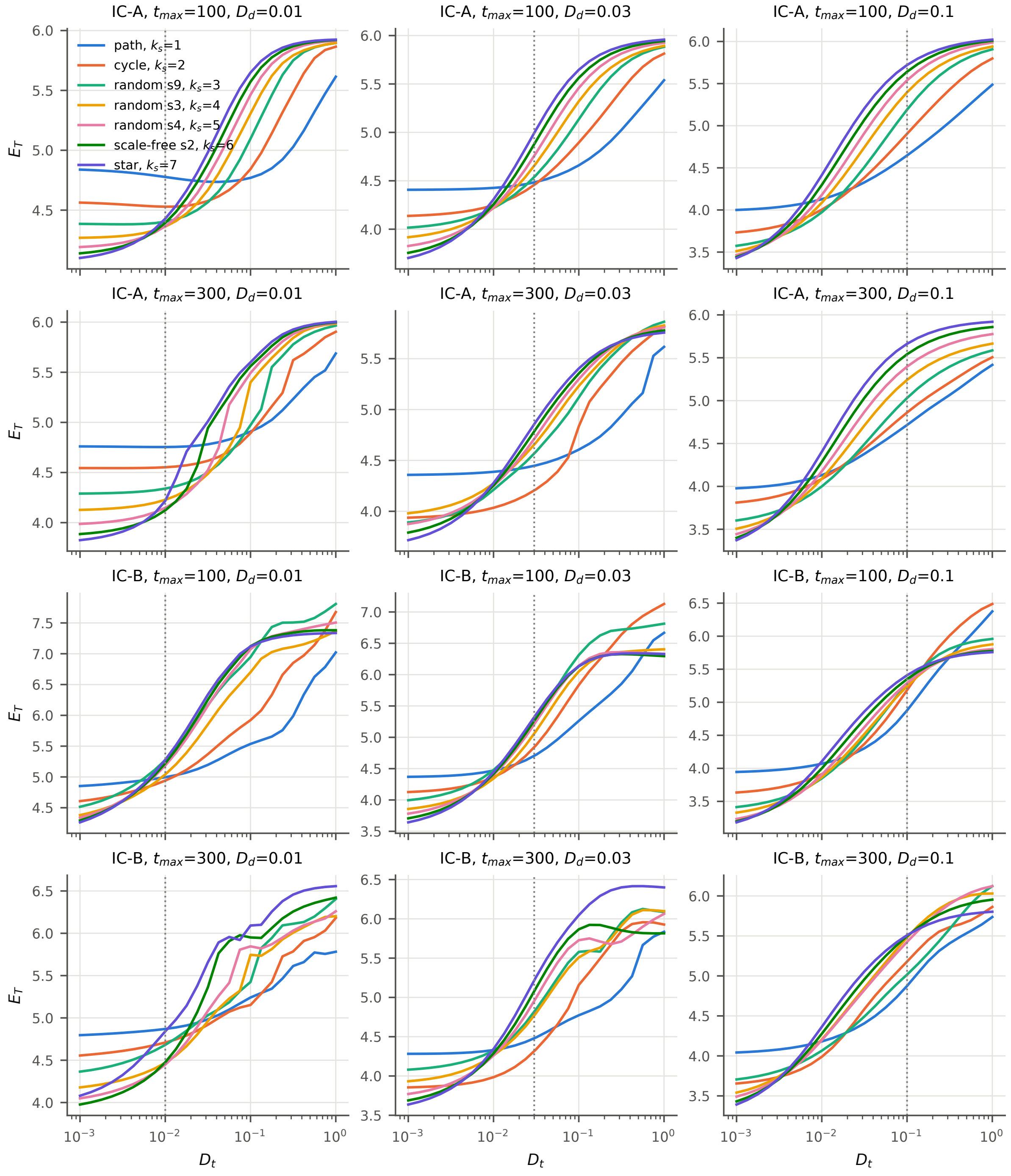


**Figure S4:** Efficiency of targeted releases against the parasitoid dispersal rate without transit cost, for every condition of Experiment 9. Rows are the initial condition and the horizon, columns the psyllid dispersal rate $D_d$. Lines are the seven focal graphs, labelled by the degree $k_s$ of the outbreak patch, and dotted lines mark $D_t = D_d$.

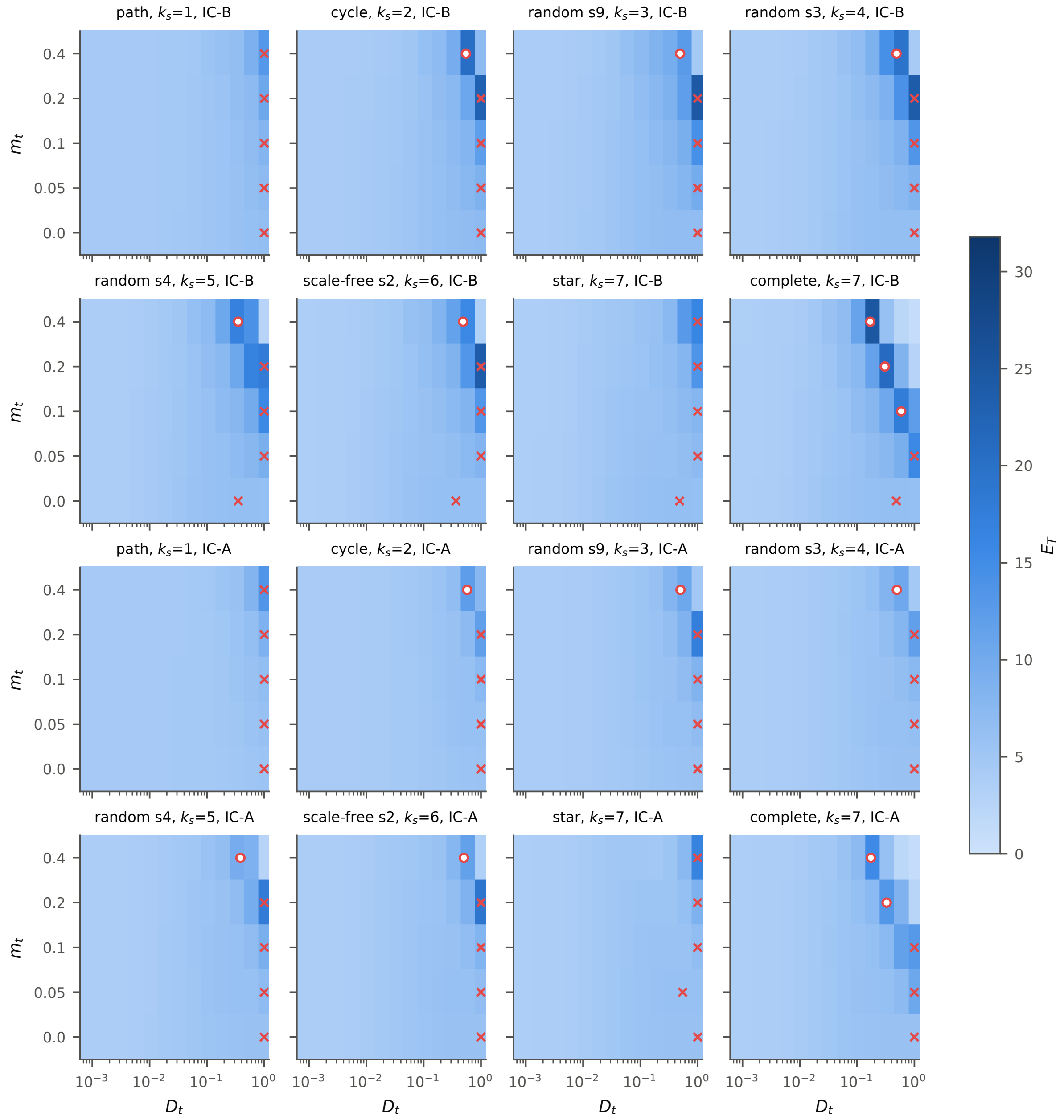


**Figure S5:** Efficiency of targeted releases on the plane ($D_t$, $m_t$) for the eight graphs of Experiment 10a under IC-B (top two rows) and IC-A (bottom two rows), with $D_d = 0.03$, $r_3 = -0.3$ and $m_d = 0$. Open circles, interior maxima with prominence above 5%; crosses, maxima at the edge of the grid or with lower prominence.

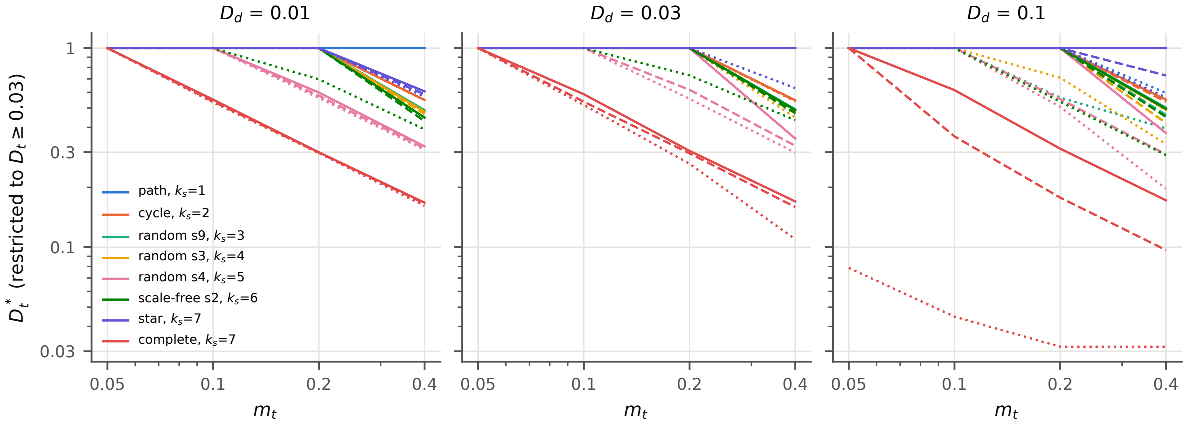


**Figure S6:** Optimal parasitoid dispersal rate $D_t^*$ against the parasitoid transit mortality $m_t$ when psyllids also die in transit (Experiment 10b, IC-B, $r_3 = -0.3$). Columns are the psyllid dispersal rate $D_d$, colours the eight graphs, and line styles the psyllid transit mortality: $m_d = 0$ (solid), 0.1 (dashed) and 0.2 (dotted). The optimum was searched for $D_t \geq 0.03$ only, so values of 1 are at the upper edge and values of 0.03 at the lower edge of the reduced grid.
